# A developmental brain-wide screen identifies retrosplenial cortex as a key player in the emergence of persistent memory

**DOI:** 10.1101/2024.01.07.574554

**Authors:** Benita Jin, Michael W. Gongwer, Brian P. Kearney, Alfonso Darmawan, Lilit Ohanian, Lucinda Holden-Wingate, Bryan Le, Yuka Nakayama, Sophia A. Rueda Mora, Laura A. DeNardo

**Affiliations:** Department of Physiology, University of California Los Angeles, Los Angeles, CA 90095; Molecular Cellular and Integrative Physiology Interdepartmental Program, University of California Los Angeles, Los Angeles, CA 90095; Neuroscience Interdepartmental Program, University of California Los Angeles, Los Angeles, CA 90095; Medical Scientist Training Program, University of California Los Angeles, Los Angeles, CA 90095; Department of Neurobiology, University of California Los Angeles, Los Angeles, CA 90095

## Abstract

Memories formed early in life are short-lived while those formed later persist. Recent work revealed that infant memories are stored in a latent state. But why they fail to be retrieved is poorly understood. Here we investigated brain-wide circuit mechanisms underlying infantile amnesia. We performed a screen that combined contextual fear conditioning, activity-dependent neuronal tagging at different postnatal ages, tissue clearing and light sheet microscopy. We observed striking developmental changes in regional activity patterns between infant, juvenile, and adult mice, including changes in the retrosplenial cortex (RSP) that aligned with the emergence of persistent memory. We then performed a series of targeted investigations of RSP structure and function across development. Chronic chemogenetic reactivation of tagged RSP ensembles during the week after learning enhanced memory in adults and juveniles, but not in infants. However, after 33 days, reactivating infant-tagged RSP ensembles recovered forgotten memories. Changes in the developmental functions of RSP memory ensembles were accompanied by changes in dendritic spine density and the likelihood that those ensembles could be reactivated by contextual cues. These studies show that RSP ensembles store latent infant memories, reveal the time course of RSP functional maturation, and suggest that immature RSP functional networks contribute to infantile amnesia.

## Introduction

Infantile amnesia describes a process by which episodic memories formed early in life are rapidly forgotten^1–3^. In contrast, memories formed later can last for decades. In humans, memories become persistent around age five^4^, but infantile amnesia is not a uniquely human phenomenon^5,6^. In rodents, episodic memories become persistent during the fourth postnatal week^7–11^. Although early-formed memories cannot be consciously recalled, they can influence later experiences by promoting the maturation of learning systems and altering mechanisms of later memory encoding^12–14^. Their continuing influence suggests that early-formed memories leave an enduring trace in the brain.

Several studies show that infant memories are not erased but rather become inaccessible. Forgotten memories leave a molecular trace in the amygdala^15^. Providing strong reminders of the original memory^13,16,17^ or optogenetically reactivating infant memory ensembles in the hippocampus^10^ can recover lost infant memories. Importantly, early adversity such as neglect or abuse can accelerate the development of emotional memory systems, ending the period of infantile amnesia prematurely^18,19^. Early adversity is a major risk factor for developing mental health disorders, suggesting that infantile amnesia may actually protect against the emergence of problematic behaviors^7,20^. Despite their importance for brain and behavioral development, the processes by which infant memories are stored and why they remain inaccessible remain poorly understood.

In adults, initial memory encoding in the hippocampus is followed by a period of consolidation, during which episodic memories come to be stored in distributed networks across the brain^21^. Memory consolidation and retrieval involve coordinated activity of numerous regions, including the prefrontal cortex, temporal cortical association areas, medial and anterior thalamic nuclei, and retrosplenial cortex (RSP)^21–23^. Connectivity among these regions continues to refine throughout postnatal development until early adulthood^24–26^. It is therefore possible that prolonged cortical development contributes to infantile amnesia. However, little is known about the functional maturation of key cortical areas. To date, mechanistic studies of infantile amnesia have focused on the hippocampus, showing that infantile amnesia is associated with naturally elevated hippocampal neurogenesis^9^, immature hippocampal synapses^13^, and immature hippocampal inhibition. As such, our understanding of which aspects of brain circuit development are associated with the emergence of persistent memories remains limited.

Here, we set out to investigate why infant memories become inaccessible. By performing contextual fear conditioning, whole-brain activity-dependent neuronal tagging^27–29^ at different postnatal ages, tissue clearing, light sheet fluorescence microscopy, and high-throughput computational analyses^30^, we identified key brain regions whose activation patterns change significantly during memory retrieval across development. The RSP emerged as a key region whose functional maturation closely aligns with changes in memory persistence. In targeted functional studies, we discovered that chronically reactivating RSP memory ensembles for a week after learning ensembles enhanced memory in adults and juveniles but not in infants. However, after 33 days, reactivation of infant-tagged RSP ensembles recovered infant memories. These developmental changes in RSP function were accompanied by enhanced RSP ensemble stability in adults and juveniles compared to infants. Tagged RSP neurons undergo significant synaptic pruning from infancy to adulthood and RSP neurons undergo learning-induced structural plasticity selectively in adults. Together, our findings demonstrate that RSP ensembles store latent infant memories but fail to be reactivated by salient contextual cues. Our studies also reveal the timeline of RSP functional maturation and indicate that low ensemble stability combined with developmental synaptic pruning contribute to infantile amnesia.

## Results

### A brain-wide screen to identify neural signatures of infantile amnesia

Using an established protocol for modeling infantile amnesia^8–10^, we performed contextual fear conditioning (CFC) in mice when they were infants (postnatal day (P) 17), juveniles (P25) or adults (P60). In separate cohorts of mice, we tested the strength of their fear memory either 1 day (d) or 7d after CFC by placing mice in the original training context and measuring freezing behavior (Figure 1A). While mice of all ages exhibited robust freezing during CFC and the 1-day (1d) memory retrieval test, mice trained as infants froze significantly less than adults and juveniles during a 7d memory retrieval test (Figure 1B).

**Figure 1.**
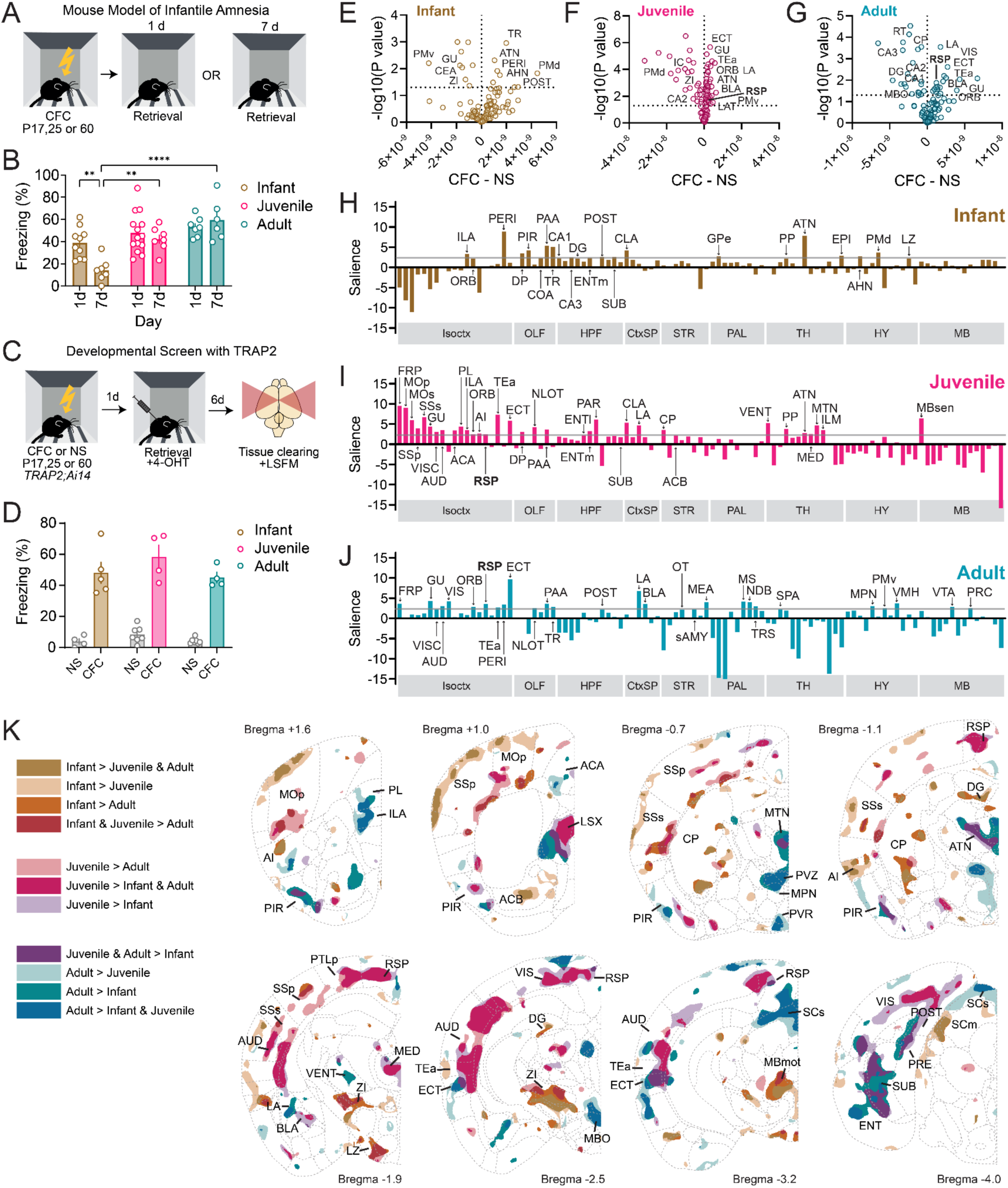
A developmental screen to identify neural signatures of infantile amnesia. (A) Infant, juvenile, and adult mice underwent contextual fear conditioning and then memory testing 1 or 7 days later. (B) Percent time freezing in infant, juvenile and adult wildtype mice during memory retrieval tests occurring 1 or 7 days after CFC (F_age_(2,45)=15.89, P<0.0001, F_day_(1,45)=3.56, P=0.07, F_int_(2,45)=4.21, P=0.02, Infant: N=7 1d, N=10 7d; Juvenile: N=14 1d, N=8 7d; Adult: N=7 1d, N=6 7d, two-way ANOVA with Tukey post-hoc test). (C) Experimental design for a developmental screen. (D) Freezing behavior in TRAP mice used in the screen. Infant: N=4 NS, N=5 CFC; P25: N=7 NS, N=4 CFC; P60: N=7 NS, N=4 CFC. (E-G) Volcano plots of unpaired two-tailed t-tests comparing learning indices between CFC and NS mice at infant (E), juvenile (F), and adult (G) timepoints. Infant: N=4 NS, N=5 CFC; P25: N=7 NS, N=4 CFC; P60: N=7 NS, N=4 CFC. Horizontal dashed line corresponds to p<0.05. (H-J) Task PLS analysis of regional learning indices in infant (H), juvenile (I), and adult (J) mice. Gray line indicates threshold value of 2. (K) Voxel-based analysis comparing learning indices in infant, juvenile, and adult CFC brains. Values from each voxel underwent one-way ANOVA with Tukey’s multiple comparisons test (Infant: N=5; Juvenile: N=4; Adult: N=4). Colored voxels indicate p<0.05 in Tukey’s test for the comparisons denoted in the legend. See Table 1 for abbreviations and Tables 2–4 for detailed statistics. Graphs show mean ± SEM. *P< 0.05, **P<0.01, ***P<0.001.

In our developmental screen, we combined activity-dependent tagging, tissue clearing, and light sheet fluorescence microscopy to screen for developmental changes in brain-wide cellular activation patterns during memory retrieval (Figure 1C). We labeled activated neuronal populations using Targeted Recombination in Active Populations (TRAP2: *Fos^icre-2A-ERT2^*)^28,29^, in which the Fos promoter drives expression of tamoxifen-inducible Cre recombinase. We crossed TRAP2 to the Ai14 Cre reporter line^31^ so all TRAPed neurons could be labeled with a red fluorophore. We performed CFC in infant, juvenile or adult *TRAP2;Ai14* mice alongside age-matched non-shocked control mice (NS), which were placed in the chamber but did not receive shocks. One day later, we TRAPed memory-activated neurons by pairing re-exposure to the training context with a tamoxifen injection. We chose to TRAP 1d instead of 7d memory retrieval because infant mice still freeze 1d after CFC (Figure 1D). This allowed us to relate neuronal activation patterns to freezing behavior while comparing the neural signatures underlying retrieval of infant memories that would fade vs. juvenile and adult memories that would persist. A week after TRAPing we perfused mice, immunostained for tdTomato and rendered brains transparent using the Adipo-Clear protocol^32^. We then imaged TRAPed cells in intact brain hemispheres using light sheet fluorescence microscopy and automatically quantified the number of TRAPed cells per brain region. After calculating a “learning index” to normalize CFC brains to age-matched NS controls (see methods), we compared the resulting patterns of neuronal activation across age groups.

First, we generated volcano plots to identify differentially activated brain regions between CFC and NS mice at each age (Figure 1E-G). In line with the onset of memory persistence, we observed similar activation patterns in CFC juvenile and adult brains but a unique pattern in infants. In juveniles and adults, CFC increased activation of association cortices including RSP, temporal association areas (TEa), and ectorhinal cortex (ECT). We also observed this effect in amygdala subregions including lateral amygdala (LA) and basolateral amygdala (BLA). In infant mice, however, CFC primarily increased activation of subcortical regions such as the dorsal premammillary nucleus (PMd), anterior hypothalamic nucleus (AHN), postpiriform transition area (TR), and anterior group of the dorsal thalamus (ATN).

To better understand which brain regions contribute to the behavioral expression of fear memories across development, we conducted a partial least-squares (PLS) analysis^33^ of TRAPed cells across brain regions. These analyses identify latent variables that maximally differentiate high versus low freezing levels during 1d memory retrieval. Bootstrap ratios (or saliences) from the PLS analysis were used to determine the extent that individual regional activation (based on numbers of TRAPed cells) contributed to the ability to distinguish freezing levels among mice within each age group. We identified regions contributing to significant contrasts between CFC and NS brains by thresholding salience values at 2, corresponding to P=0.05 (Figure 1H–J). In infant mice, we observed high salience in olfactory (OLF) and hippocampal regions (HPF), in some thalamic nuclei (TH) and in the hypothalamus (HY) (Figure 1H). In juveniles, we observed high salience values throughout the isocortex, in several thalamic nuclei, and in retrohippocampal and olfactory areas (Figure 1I). Salient regions in adults were dispersed throughout the brain, with salient regions in the hypothalamus and midbrain that were distinct from the other ages (Figure 1J). Consistent with our region-based analyses, we found overlap in the salient regions of juvenile and adult mice in association cortices such as RSP, TEa, and ECT, which we did not observe in infant mice.

We next sought to describe within-region variations in age-specific memory activation. To achieve this, we segmented brains into individual 10 μm voxels, with each voxel representing the cell density of the surrounding ∼500 μm. We normalized each voxel to that of age-matched NS controls by calculating the learning index as before, then performed a one-way ANOVA for each voxel in CFC brains across ages (Figure 1K). A large number of voxels throughout the brain were significantly modulated by age, often concentrated in particular brain regions. We first focused on voxels that had a significantly higher learning index in juvenile and adult brains compared to infant brains (shown in dark purple in Figure 1K), as this aligns with the onset of memory persistence. We observed this effect in the piriform cortex (PIR), midline thalamic nuclei (MTN), ATN, ECT, TEa, and subicular subregions (SUB, PRE, POST). We also observed this in subregions of RSP. In contrast, we found that voxels in the hypothalamic zona incerta (ZI) and motor areas of the midbrain (MBmot) had a higher learning index in infant mice compared to older animals. Juvenile mice had a uniquely high learning index in several posterior cortical regions and the lateral septum (LSX). Adult mice had high learning indices in prelimbic and infralimbic cortices (PL & ILA) as well as the hypothalamic periventricular zone (PVZ), medial preoptic nucleus (MPN), and mammillary body (MBO).

Several similar patterns emerged in voxel-based comparisons between CFC and NS animals at each age (Figures S1-S3). Compared to NS controls, CFC infants had higher learning indices in the hypothalamus. In contrast, CFC juveniles and adults had higher learning indices in subregions of the cortex, including RSP, amygdala and hippocampal formation compared to age-matched NS mice. An age-dependent shift in the distribution of 1d memory recall ensembles again appears between infants and juveniles, indicating that changes in fear memory networks align with the developmental shift in memory persistence.

Overall, our developmental screen provided convergent evidence that a network of regions known to play a key role in memory consolidation and long-term storage in adults –including RSP, TEa, ECT, LA, and BLA– are not recruited to the same extent in infant mice, whose memory traces are marked by a unique network of olfactory, hypothalamic, and hippocampal regions. While RSP has established roles in memory consolidation and long-term memory retrieval in adults^34–39^, its role in memory is poorly understood in the developing brain. Based on the convergent findings in our screen, we chose to further examine the role of RSP in memory retrieval and consolidation across development.

### Reactivation of TRAPed RSP ensembles shortly after learning enhances memory in an age-dependent manner

Offline activity within memory networks is thought to underlie memory consolidation^21^. A recent study revealed that re-activation of tagged RSP memory ensembles can accelerate memory consolidation^36^. We therefore hypothesized that offline reactivation of RSP memory ensembles could recover lost infant memories. To test this, we injected an AAV into RSP expressing the excitatory Cre-dependent DREADD (designer receptor exclusively activated by designer drugs) hM3Dq^40^ or a control AAV that encoded a Cre-dependent fluorophore. Two weeks later – when mice were infants, juveniles, or adults – we performed contextual fear conditioning. The following day, we TRAPed RSP neurons activated by 1d memory retrieval, thereby driving expression of hM3Dq in those cells. We then performed off-line reactivation of the TRAPed ensembles by administering the hM3Dq ligand clozapine-N-oxide (CNO, 1 mg/kg/day, intraperitoneally) daily. We administered a final dose of CNO (2 mg/kg) 30 min prior to a test retrieval session on day 7 (Figure 2A,B). Histology confirmed that hM3Dq-mCherry was expressed in a similar location within RSP and to a similar extent across age groups (Figure S4) and that hM3Dq expressing mice had significantly higher levels of activity in RSP (as assayed with Fos immunostaining) compared to controls (Figure S5).

**Figure 2.**
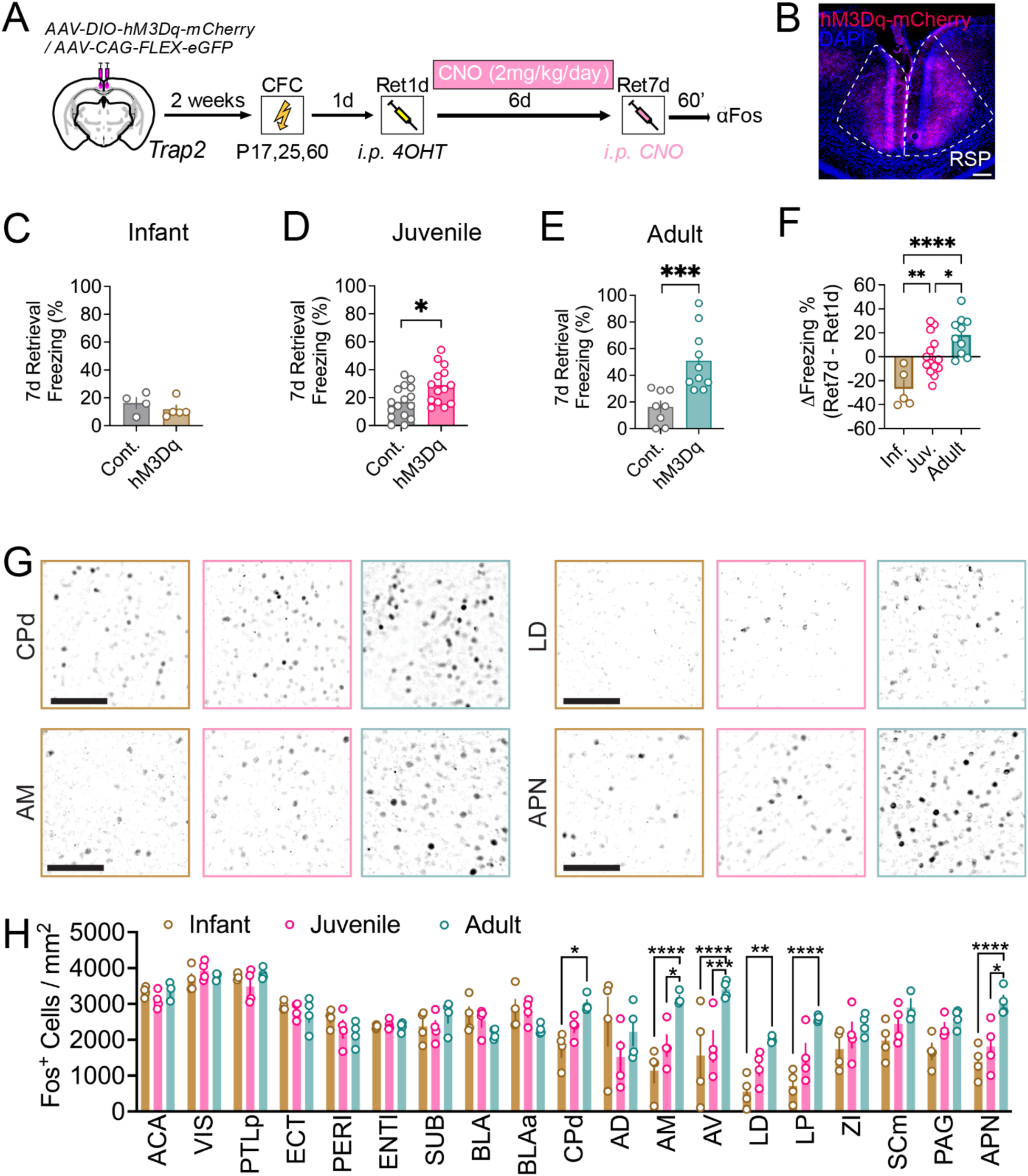
Chemogenetic reactivation of TRAPed RSC neurons increased freezing during short-term memory retrieval in an age-dependent manner. (A) Experimental design. (B) Representative coronal section showing AAV injection site in RSP. Scale bar, 125 µm. (C) Percent freezing levels at 7-day retrieval in control (grey) and hM3Dq (brown) groups for infants (P=0.36 Unpaired t test). (D) Percent freezing levels at 7-day retrieval in control (grey) and hM3Dq (magenta) groups for juveniles (P=0.01 Unpaired t test). (E) Percent freezing levels at 7-day retrieval in control (grey) and hM3Dq (teal) groups for adults (P=0.001 Unpaired t test). (F) Differences in freezing levels between 7-day and 1-day retrieval sessions for hM3Dq mice in each age group (F(2,26)=12.92, P=0.001, Infant: N=5, Juvenile: N=14, Adult: N=10, one-way ANOVA with Tukey post hoc test). (G) Representative images showing Fos immunoreactive cells in dorsal caudate putamen (CPd), anteromedial thalamus (AM), laterodorsal thalamus (LD), and anterior pretectal nucleus (APN). Scale bars, 100 µm. (H) Quantification of Fos+ cells per ROI in 19 brain regions (F_region_(18,160)=30.03, P<0.05; F_age_(2,9)=3.69, P>0.05; F_int_(36,160)=5.78, P<0.05, Mixed Effects Model; Infant: N=4; Juvenile: N=4; Adult: N=3-4). Graphs show mean ± SEM. * P<0.05, **P<0.01, ***P<0.001, ****P<0.0001. See also Figure S4 and S5. See Table 1 for abbreviations.

Chronic chemogenetic activation of TRAPed RSP ensembles shortly after learning affected fear memory retrieval in an age-dependent manner. In infant mice, both control and hM3Dq groups froze minimally during 7d retrieval, indicating that enhancing activity in RSP memory ensembles in infant mice cannot prevent infantile amnesia (Figure 2C). Juvenile hM3Dq-expressing mice froze significantly more than controls during 7d retrieval (Figure 2D). In line with previous findings^36^, RSP ensemble reactivation in adult mice significantly increased conditioned freezing during 7d retrieval compared to both the 1d retrieval session and to 7d retrieval in control mice (Figure 2E). To further compare age-specific behaviors, we computed for each hM3Dq-expressing mouse the change in freezing levels from 1d to 7d retrieval (Figure 2F). All infants froze less, juveniles had no net change in freezing, and adults froze more by the 7d test. Together, these data suggest that mature functions of RSP emerge between juvenile and adult stages, and that memories are consolidated or retrieved via a distinct mechanism in the infant brain.

To investigate the circuit-level differences underlying the distinct, age-specific effects of reactivating RSP ensembles, we sacrificed the mice 60 min after the 7d retrieval test for Fos immunostaining. We sectioned the brains, aligned individual slices to the Allen Brain Atlas and then quantified the number of Fos+ neurons across a variety of brain regions. We focused on brain regions that receive direct anatomical projects from RSP or have known functional connectivity with RSP^25,35,41–45^. Compared to adults, infants had significantly fewer Fos+ neurons in the dorsal caudate putamen (CPd), several anterior thalamic nuclei (AM, AV, LD and LP), and the anterior pretectal nucleus (APN), a midbrain motor nucleus (Figure 2G,H), suggesting that immature functional connectivity between RSP and these regions may lead to retrieval failure in infants.

### Reactivation of infant-TRAPed RSP ensembles weeks after learning can recover lost memories

We next investigated whether infant-TRAPed RSP ensembles could promote memory retrieval after development had progressed. To test this, we injected an AAV encoding Cre-dependent hM3Dq or a fluorophore control into RSP. Two weeks later, when mice were P17, we TRAPed RSP ensembles during contextual fear conditioning. 33d later, we delivered CNO before a memory retrieval test (Figure 3A,B). As before, histology confirmed that hM3Dq-mCherry was targeted to RSP and that hM3Dq-expressing mice had significantly higher levels of activity in RSP (as assayed with Fos immunostaining) compared to controls (Figure S6).

**Figure 3.**
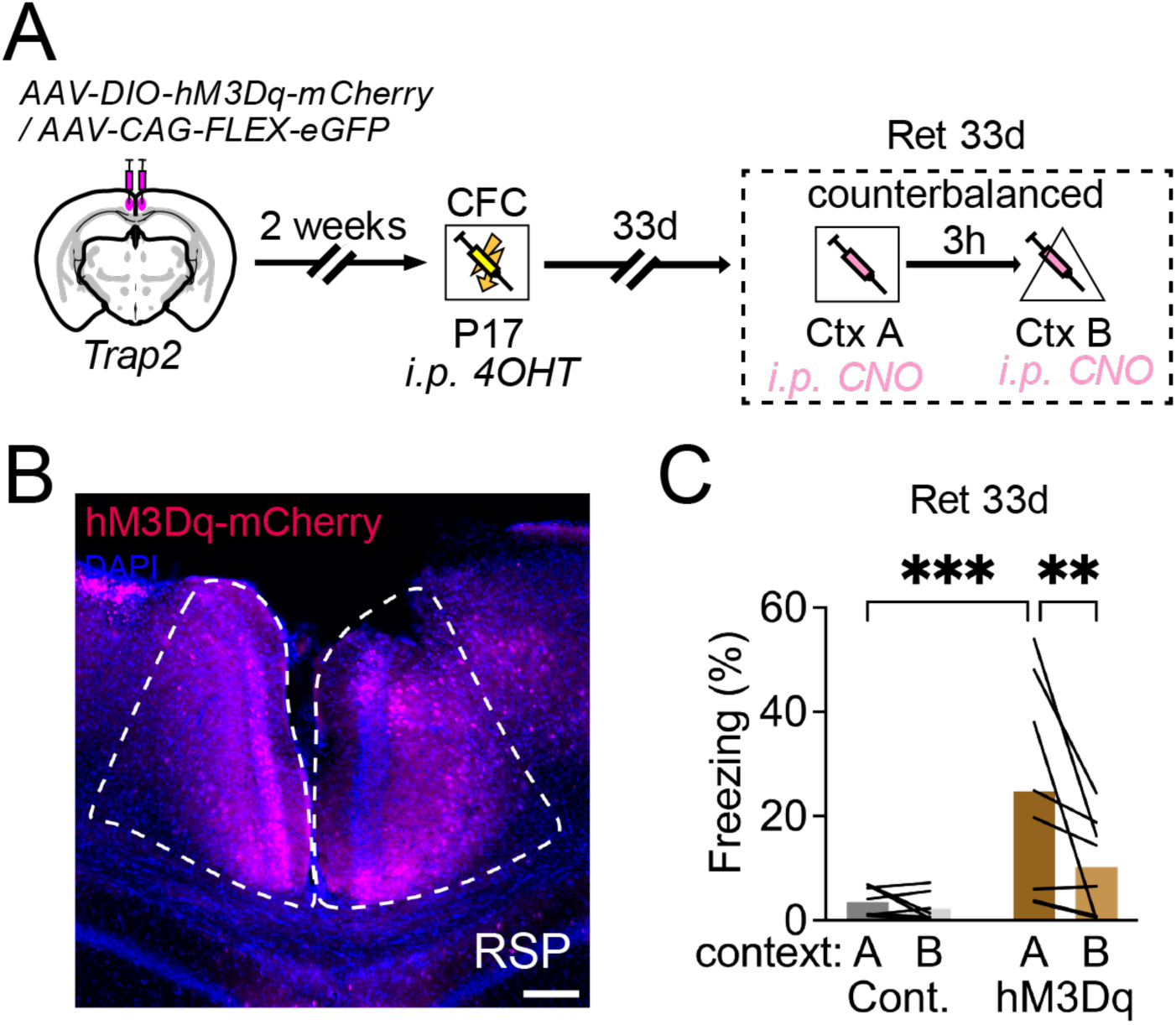
Chemogenetic reactivation of learning encoding ensembles in RSC 33d after training recovers infant memories (A) Experimental design. (B) Representative coronal section showing AAV injection site in RSP. Scale bar, 125 µm. (C) Percent freezing levels in contexts A and B during the 33-day retrieval test in control (gray) and hM3Dq (brown) mice (F_virus_(1,14)=9.06, P=0.009; F_context_(1,14)=7.41, P=0.01; F_int_(1,14)=5.23, P=0.03, Cont: N=8, hM3Dq: N=8, two-way ANOVA with Holm-Sidak posthoc test). Graphs show mean ± SEM. *P<0.05. See also Figures S6 and S7.

In this experiment, hM3Dq-expressing mice froze significantly more than controls. We observed this effect in the conditioning context, but not in an altered context that had a distinct floor, wall decorations and scent (Figure 3C). Together, these results reveal that infant RSP ensembles store latent memories and that once RSP cells and circuits develop, reactivating those ensembles can recover the stored infant memory. However, because activating TRAPed RSC cells only drove freezing in the conditioning context, our findings also indicate that the function of infant-TRAPed RSP ensembles are gated by the presence of salient contextual cues. Reactivating infant-TRAPed neurons in ACA or LSX did not enhance freezing 33d later (Figure S7), suggesting the effects we observed are specific to RSP.

### Reactivation of TRAPed RSP neurons during memory retrieval increases with age

Memory persistence has been ascribed to the reinstatement of patterns of neuronal ensemble activity that were established during initial learning^46,47^. In the hippocampus, mPFC and RSP, neuronal ensembles expressing Fos during learning or recent memory retrieval overlap with Fos+ ensembles evoked during later recall^28,48^. The extent of ensemble reactivation predicts retrieval success^5,26^. While RSP is known to play key roles in memory consolidation and remote memory retrieval^35,49^, how ensemble stability in the RSP changes across development remains unknown.

To determine to the extent to which ensembles of RSP neurons are reactivated during memory retrieval across time, we used a combination of TRAP2 and immunostaining for Fos, a marker of recently activated neurons (Figure 4A). Infant, juvenile, or adult mice were TRAPed during CFC, permanently labeling activated neurons with tdTomato. Then, following a 7d memory retrieval test in the same mice, brains were collected for Fos immunostaining (Figure 4B). In RSP, we observed no differences in the numbers of TRAPed or Fos+ cells after CFC (Figure 4C-E). However, compared to infants, juvenile and adult mice had significantly higher levels of reactivated neurons in RSP (Figure 4F). These findings indicate that greater memory ensemble stability in RSP aligns with the emergence of persistent episodic memories.

**Figure 4.**
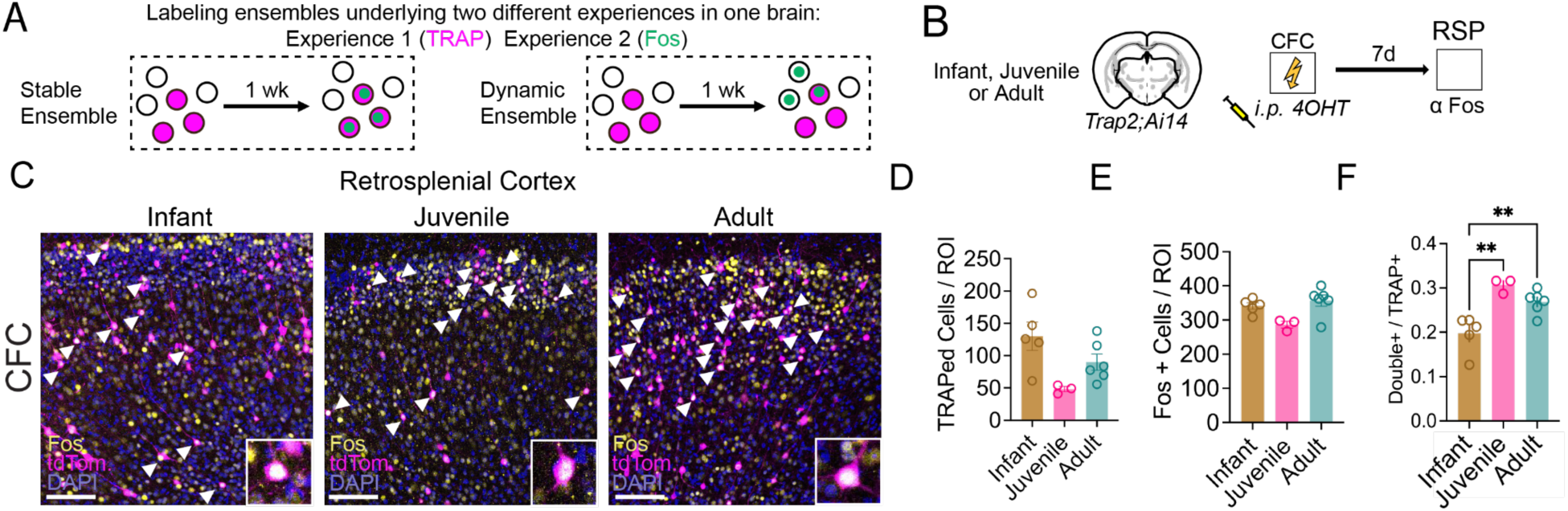
Selectively increased reactivation of memory-encoding ensembles in RSP in juveniles and adults. (A) Illustration of stable vs. dynamic ensembles in a TRAP/Fos overlay assay. (B) Experiment design for activity-dependent tagging of ensembles at CFC and Fos staining following 7-day retrieval. (C) Representative examples of TRAPed (magenta) and Fos+ cells (green) in RSP. Double-labeled cells are in white. (D) CFC-TRAPed RSP cells (F(2,11)=12.88, P=0.0013, one-way ANOVA with Tukey post-hoc correction; Infant: N=4 mice, Juvenile: N=8 mice, Adult: N=7 mice; 3 coronal sections per mouse). (E) Fos+ cells in RSP after 7d retrieval (F(2,16)=2.11, P=0.15, one-way ANOVA with Tukey post-hoc correction; Infant: N=4 mice, Juvenile: N=8 mice, Adult: N=7 mice; 3 coronal sections per mouse). (F) Double-labeled neurons in RSP after fear learning and 7d retrieval (F(2,11)=12.88, P=0.0013, one-way ANOVA with Tukey post-hoc correction; Infant: N=4 mice, Juvenile: N=4 mice, Adult: N=4 mice; 3 coronal sections per mouse). Graphs show mean ± SEM. *P <0.05. Scale bar, 100 µm.

### Selective learning-dependent structural plasticity in memory-encoding RSP neurons in adults

We next investigated whether synaptic mechanisms of learning in RSP may change across development. Dendritic spines are key sites of sites of excitatory synaptic contact. Structural plasticity of dendritic spines in RSP has been linked to fear learning and memory^50,51^. Dendritic spine turnover and clustering in the adult RSP correlates with the strength of fear learning and memory^51^. To determine whether developing RSP ensembles undergo learning-induced structural plasticity, we fear conditioned TRAP2;Morf3 mice and TRAPed memory encoding ensembles (Figure 5A). In the presence of tamoxifen, TRAP2;Morf3 mice express a V5 tag in a subset of activated neurons, resulting in very bright, sparse labeling that enables detailed morphological analysis, including quantification of dendritic spines (Figure 5B). In NS control mice, RSP dendritic spine density decreased steadily across development (Figure 5C), suggesting there is ongoing synaptic pruning between infancy and adulthood. In fear conditioned mice, infants had slightly lower spine density compared to NS controls, juveniles had similar spine density to NS controls and adults had significantly higher spine density than NS controls (Figure 5C). Since we did not observe structural plasticity in juveniles, which have persistent memory, these findings suggest that RSP structural plasticity may not be critical for the emergence of persistent memory. However, these data suggest that developmental pruning in RSP may contribute to the failure to retrieve infant memories^52^.

**Figure 5.**
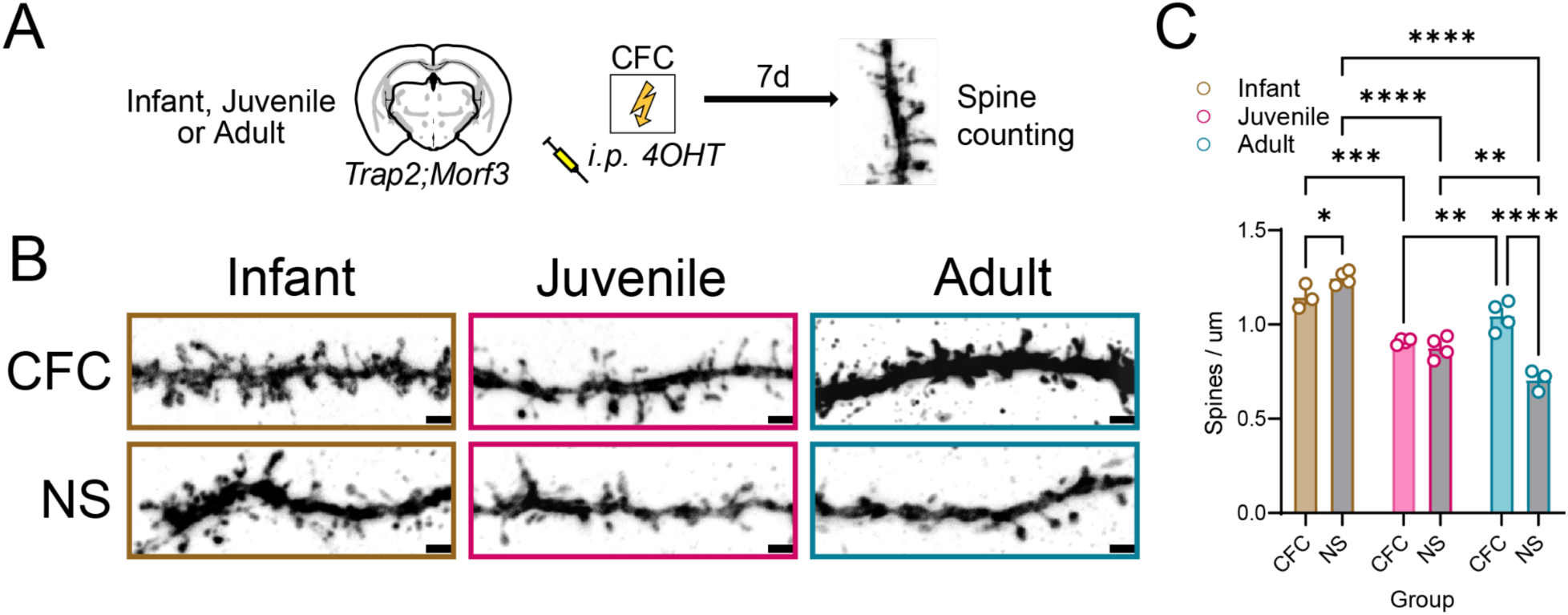
Selective learning-dependent structural plasticity in memory-encoding RSP neurons in adults. (A) Experiment design for activity-dependent tagging of dendritic spines during learning. (B) Representative examples of TRAPed spines during CFC or NS learning in infants (brown), juveniles (magenta), or adults (teal). (C) Fear learning increases TRAPed RSP spine density with age (colored bars, F_group_(1,16)=14.62, P=0.0015, two-way ANOVA with Tukey post-hoc correction; Infant: N=4 mice, Juvenile: N=4 mice, Adult: N=4 mice; 6-8 dendritic segments analyzed per mouse). Non-shock controls show a gradual decrease in spine density with age (gray bars, F_age_(2,16)=75.79, P<0.0001, two-way ANOVA with Tukey post-hoc correction; Infant: N=4 mice, Juvenile: N=4, Adult: N=3 mice; 6-8 dendritic segments analyzed per mouse). F_interaction_(2,16)=29.29, P<0.0001, two-way ANOVA with Tukey post-hoc correction. Graphs show mean ± SEM. *P <0.05, **P <0.01, ***P <0.001, ****P <0.0001. Scale bars, 2µm.

## Discussion

Even though very early experiences cannot be explicitly recalled, they can profoundly impact the rest of our life^5,7,14^. Recent work shed light on this paradox, showing that infant memories are not erased but rather inaccessible^10,13,15–17^. But why infant memories become inaccessible is poorly understood. To investigate this, we designed a brain-wide screen to identify developmental changes in memory retrieval circuits. We found a number of changes in regional activation patterns that aligned with the onset of persistent memory, including greater memory-induced activation in RSP (Figure 1). We then defined the time course of RSP functional maturation. We found that chronically reactivating TRAPed RSP ensembles during the week after CFC enhanced memory in adults and juveniles, but not in infant mice (Figure 2). On the other hand, activating infant-TRAPed RSP ensembles more than a month after CFC rescued infant fear memories (Figure 3). Changes in the developmental functions of RSP memory ensembles were accompanied by increases in the likelihood that juvenile and adult RSP memory ensembles would be reactivated by contextual cues (Figure 4), developmental spine pruning in RSP, and learning-dependent structural plasticity that was unique to adults (Figure 5). Together these findings reveal that learning-activated RSP ensembles encode latent infant memories across time. However, salient sensory cues do not activate RSP cells that encode infant memories - perhaps due to synapse loss - so retrieval does not occur. Furthermore, our studies delineate the functional maturation of RSP as it relates to memory encoding and retrieval.

Current models of infantile amnesia center on hippocampal maturation. In infants, learning promotes hippocampal synaptic maturation via signaling pathways that are involved in critical periods in other brain regions, suggesting that the infant hippocampus is in a critical period for learning systems^13^. These studies led the authors to propose the retrieval hypothesis, in which an immature hippocampus stores information in a latent form such that it fails to be retrieved without extensive reminders. In this model, learning promotes hippocampal maturation and drives transitions through sequential critical periods. In each critical period, the hippocampus becomes better at storing information such that it can be actively retrieved later^14^. Other studies showed that in the infant brain, naturally high levels of hippocampal neurogenesis promote forgetting; but infant memories are stored in a dormant state in hippocampal ensembles^9,10^. This led the authors to propose a (non-mutually exclusive) neurogenic hypothesis of infantile amnesia, in which neurogenesis-induced circuit remodeling underlies infantile amnesia^5,53^.

Our findings build upon this work, revealing brain wide changes in cellular activation during memory retrieval across development. The neural mechanisms of memory retrieval are well studied in adults but remain poorly understood in the developing brain. At the time point we examined (1d after CFC), memory consolidation was underway, so the data presented in Figure 1 may provide insight into the circuits participating in memory consolidation as well as memory retrieval. Our whole brain analyses revealed that juveniles and adults had marked learning-dependent activation of RSP, TEa, ECT, and LA (Figure 1F,G,I,J), areas with known roles in memory retrieval and consolidation^35,38,54^. On the other hand, infants had more activation in hypothalamic nuclei including PMd and ZI (Figure 1E,H). In the adult brain, these regions are associated with spatial escape and fear learning and memory, respectively, but their developmental functions are poorly understood^55,56^. Even though juveniles and adults had similar freezing levels across time, we also observed distinct cellular activation patterns between these two age groups, including in the prefrontal cortex and midbrain (Figure 1K). These may be important for increases in memory complexity or specificity that emerge later in development, after memories become persistent. Together, our findings suggest that juvenile and adult fear memories likely have unique circuit mechanisms of memory consolidation, storage and retrieval. Future studies can manipulate activity in the hypothalamic regions we identified, for example, to determine their role in fear learning and memory in the developing brain.

Our findings further suggest that, in addition to the hippocampus, the immature RSP also contributes to the retrieval failure of infant memories. In adults, both in rodents and humans, RSP plays a key role in learning^37,60,61^, consolidation^34,49,62,63^ and recent and remote memory retrieval^59,64–66^, but how and when RSP functionally matures was largely unknown. We provide novel insights into the RSP circuit mechanisms underlying memory retrieval and consolidation in adults and align functional maturation of RSP with its changing role in memory. One caveat of the chemogenetic studies presented in Figure 2 is that we activated TRAPed neurons offline and immediately before memory retrieval. Therefore, we cannot distinguish the role of RSP in memory consolidation vs. retrieval. Still, our results suggest that while activating TRAPed RSP neurons enhances memory in juveniles and adults, these functions are absent in infants. Our studies further suggest that developmental pruning of key RSP connections contributes to memory retrieval failure. The RSP is densely interconnected with sensory cortices, with limbic regions that are important for memory encoding, and with anterior thalamic nuclei and cortical regions important for long-term memory storage and retrieval^21,39,43,44,55,67–74^. In developing rats, bidirectional connections linking RSP to the hippocampal formation reach adult-like densities by P12^25,26^. In humans, resting state functional connectivity between RSP and the prefrontal cortex increases between adolescence and early adulthood while RSP-hippocampal functional connectivity matures earlier^75^. Future studies can investigate which key connections are pruned.

It remains to be determined how an immature hippocampus may influence the ability of the infant RSP to encode, consolidate, and retrieve memories. RSP serves as an input structure to the hippocampus, providing important spatial cues. RSP also serves as a target of hippocampal output during systems consolidation^57^. In adults, the transfer of information from the hippocampus to RSP is critical for the formation of long-term memories in the RSP^58–60^ and for subsequent memory retrieval^57,61,62^. In infants, the immature hippocampus could lead to weaker memory representations in RSP as compared to adults. But we show that directly stimulating infant-TRAPed RSP ensembles can recover the infant memory, suggesting that some post-learning hippocampal-RSP information transfer has taken place. Further, our analyses indicate that in infants, weaker functional connectivity between RSP and anterior thalamic nuclei and midbrain motor areas may underlie retrieval failure. Future studies can examine how RSP connections with the hippocampus, anterior thalamic nuclei, and midbrain influence the emergence of persistent memory.

Early life adversity like chronic stress is a major risk factor for developing mental health disorders. Early life adversity can lead to premature emergence of persistent memories, suggesting that accelerated development of memory systems may be linked to poor mental health outcomes^7,18–20^. In aging models of neurodegeneration, in which subjects display a similar level of forgetting as in infantile amnesia, memory impairment corresponds to the degeneration of memory circuits that include the RSP^63,64^. By illustrating how memory retrieval networks change across development, our work builds a foundation for understanding how adverse rearing conditions may influence neurodevelopmental trajectories, leading to problematic behaviors in adulthood. Further, revealing the organization and function of memory networks in animals with infantile amnesia may provide mechanistic insight into memory disorders associated with forgetting^65^.

## Acknowledgments

We thank Carlos Portera-Cailliau and Scott Wilke for feedback on the manuscript. This publication was supported by a NARSAD Young Investigator Award from BBRF (L.A.D.), Whitehall Foundation Grant (L.A.D.), Klingenstein-Simons Fellowship in Neuroscience (L.A.D.), R01MH127214 (L.A.D), T32 MH073526 (B.J.), Jessamine Hilliard Neurobiology Graduate Student Grant Program (B.J) and UCLA Dissertation Year Fellowship (B.J.). Confocal imaging was performed in the UCLA Broad Center for Regenerative Medicine and Stem Cell Research microscopy core.

## Author Contributions

Conception, B.J. and L.A.D.; Methodology, B.J., M.W.G., B.P.K., L.O., L.H.W., B.W., A.D., Y.N. S.A.R.M., and L.A.D.; Investigation, B.J.,; Writing – Original Draft, B.J. and L.A.D.; Writing – Review & Editing, B.J., M.W.G., L.A.D.; Funding Acquisition, B.J. and L.A.D.

## STAR Methods

### Resource availability

Further information and request for resources and reagents should be directed to and will be fulfilled by the lead contact, Laura A. DeNardo

### Materials availability

This study did not generate new unique reagents.

### Data and code availability

Whole brain TRAP data are presented in Table 2–4 and previously published code for DeepCOUNT can be found in the DeNardo lab GitHub database (https://github.com/DeNardoLab). Any additional information required to reanalyze the data reported in this paper is available from the lead contact upon request.

### Experimental model and study participant details Animals

Female and male C57B16/J mice (JAX Stock No. 000664) or TRAP2;Ai14 mice (JAX Stock Nos. 030323 and 007914) were group housed (2–5 per cage) and kept on a 12 hour light cycle (lights on 7am-7pm). All animal procedures followed animal care guidelines approved by the University of California, Los Angeles Chancellor’s Animal Research Committee.

## Method Details

### Surgery

Animals underwent surgery at P3, P9 or P46. Depending on the experiment, mice were anesthetized through hypothermia (P3 mice), or induced in 5% isoflurane in oxygen until loss of righting reflex (P9 or older) and transferred to a stereotaxic apparatus with rubber ear bars. P9+ mice were maintained under 2% isoflurane in oxygen, and warmed with a circulating water heating pad throughout surgery and eye gel was applied to the animal’s eyes. The mouse’s head was shaved and prepped with three scrubs of alternating betadine and then 70% ethanol. Following a small skin incision, a dental drill was used to drill through the skulls above the injection targets. A syringe pump (Kopf, 693A) with a Hamilton syringe was used for injections delivered at a rate 100nL/min. The syringe was left in the brain for 1 or 7 minutes (min) following injection for P3 and P9+ mice respectively. For DREADD studies, a small volume of AAV8-hSyn-DIO-hM3D(Gq)-mCherry virus (Addgene 44361, final titer 2 x 10^12^ pp per mL) was injected bilaterally into RSC (AP: −2.7, ML: ± 0.3, DV: 1.0). At P3 and P9, this coordinate was scaled using Neurostar StereoDrive software (Neurostar) which corrects for head size, orientation, and tilt. To compensate for age-dependent changes in cortical volume, we made proportional adjustments to the volume of injected virus based on previous developmental studies^84^: 0.15 µL for P3, 0.2 µL for P9, and 0.4 µL for P46 mice. Control mice received injections of age-matched volumes of AAV1-pCAG-FLEX-eGFP-WPRE (Addgene 51502, final titer 1 x 10^13^ pp per mL) into RSC. For pain management mice received 5 mg/kg carprofen diluted in 0.9% saline subcutaneously. Mice received one carprofen injection 30 min prior to surgery and daily injections for two days following surgery. Following confirmatory histology, samples with mistargeted injection sites were excluded from analysis.

### 4-Hydroxytamoxifen Preparation

4-hydroxytamoxifen (4-OHT; Sigma, Cat# H6278) was dissolved at 20 mg/mL in ethanol by shaking at 37°C for 15 min and was then aliquoted and stored at –20°C for up to several weeks. Before use, 4-OHT was redissolved in ethanol by shaking at 37°C for 15 min, a 1:4 mixture of castor oil:sunflower seed oil (Sigma, Cat #s 259853 and S5007) was added to give a final concentration of 10 mg/mL 4-OHT, and the ethanol was evaporated by vacuum under centrifugation. The final 10 mg/mL 4-OHT solutions were always used on the day they were prepared. All injections were delivered intraperitoneally (i.p.).

### Behavioral assays

Mice were placed in an operant chamber with a shock floor. For context fear conditioning, mice received 5 1 second, 0.5 mA foot shocks separated by 1 min. On retrieval days, mice were placed in the conditioning chamber for 5 min. For TRAP experiments, mice were injected with 4-OHT solution immediately following the retrieval session. For quantifying MORF+ cells, learning sessions were for two days, with 3 shocks on day 1 and 2 shocks on day 2. Point-tracking of behavioral videos was performed in DeepLabCut^66^. Locomotion and freezing behavior was analyzed using BehaviorDEPOT^67^.

### Brain Clearing

Two weeks after TRAPing, mouse brains were collected and processed based on the published Adipo-Clear protocol^32^ with slight modifications. Mice were perfused intracardially with 20mL of PBS (Gibco) followed by 4% paraformaldehyde (PFA, Electron Microscopy Sciences) on ice. Brains were hemisected approximately 1 mm past midline and postfixed overnight in 4% PFA at 4°C. The following day, samples were dehydrated with a gradient of methanol (MeOH, Fisher Scientific):B1n buffer (1:1,000 Triton X-100, 2% w/v glycine, 1:10,000 NaOH 10N, 0.02% sodium azide) for 1 hour for each step (20%, 40%, 60%, 80%) on a nutator (VWR). Samples were then washed with 100% MeOH 2x for 1 hour each and then incubated in a 2:1 dicholoromethane (DCM):MeOH solution overnight. The following day, two washes of 1 hour in 100% DCM were performed followed by three washes of 100% MeOH for 30 min, 45 min then 1 hour. Samples were bleached for 4 hours in 5:1 H2O2/MeOH buffer. A cascade of MeOH/B1n washes (80%, 60%, 40%, 20%) for 30 min each rehydrated the samples followed by a 1 hour wash in B1n buffer. 5%DMSO/0.3M Glycine/PTxWH permeabilized tissue for one hour and then again for 2 hour with fresh solution. Samples were washed with PTxwH for 30 min and then incubated in fresh PTxwH overnight. The following day two more PTxwH washes lasted 1 hour then 2 hours. For TRAP2 experiments, samples were incubated in primary RFP antibody (Rockland 600-401-379) at 1:300 in PTxwH shaking at 37°C for 11 days, washed in PTxwH 2×1 hour and then 2×2 hours, then for two days with at least one PTxwH change per day while shaken at 37°C. Samples were then incubated in secondary antibody AlexaFluor 647, catalog #A-31573, 1:300, ThermoFisher Scientific) for 8 days shaken at 37°C. Samples were washed in PTxwH 2×1 hour, then 2×2 hour, then 2 days with at least one PTxwH change per day while shaken at 37°C. Samples were then washed in 1x PBS twice 1×1 hour, 2×2 hours and then overnight. To dehydrate samples, a gradient of washes in MeOH:H2O (20%, 40%, 60% and 80%) were conducted for 30 min each, followed by 3×100% MeOH for 30 min, 1 hour, then 1.5 hours. Samples were incubated overnight in 2:1 DCM:MeOH on a nutator. The next day, samples were washed in 100% DCM 2×1 hour each. Samples were then cleared in 100% dibenzyl ether (DBE). DBE was changed after 4 hours. Samples were stored in DBE in a dark place at room temperature. Imaging took place at least 24 hours after clearing. Samples with poor antibody penetration following tissue clearing were excluded from analysis. To limit variation due to behavioral performance in the CFC condition, only mice with at least 35% freezing at 1-day retrieval were included in further analyses.

### Whole Brain Imaging

Brain samples were imaged on a light-sheet microscope (Ultramicroscope II, LaVision Biotec) equipped with a sCMOS camera (Andor Neo) and a 2x/0.5 NA objective lens (MVPLAPO 2x) equipped with a 6 mm working distance dipping cap. Image stacks were acquired at 0.8x optical zoom using Imspector Microscope v285 controller software. For TRAP2;Ai14 brains, we imaged using 488-nm (laser power 20%) and 640-nm (laser power 50%) lasers. The samples were scanned with a step-size of 3 µm using the continuous light-sheet scanning method with the included contrast adaptive algorithm for the 640-nm channel (20 acquisitions per plane), and without horizontal scanning for the 488-nm channel.

### DeepCOUNT Analysis

#### Image Registration & Cell Segmentation

Whole-brain TRAP counts were obtained using DeepCOUNT as previously described^30^. Briefly, whole-brain image stacks from the 640nm channel from each brain were segmented in TrailMap^68^ using the TRAP2-Ai14 trained model. Images were registered to the template brain using elastix and transformix using the 488nm autofluorescence channel. MATLAB was used to identify 3D maxima of the transformed probability map. Connected component analysis was used to reduce any maxima that consisted of multiple pixels into a single pixel per cell.

#### Cell Quantification

Regional TRAPed cell density was quantified in MATLAB (Mathworks) by counting the number of labeled pixels (i.e. cells) in each brain region, then dividing this pixel count by the total number of pixels in a region. We observed variability in the efficiency of tamoxifen-induced recombination between brains as measured by the total number of TRAPed cells across regions. To account for region- and age-specific differences in recombination efficiency, a linear regression was fitted between the total number of cells in a brain (a proxy for recombination efficiency) and cell counts in a particular region for non-shocked control brains within an age.

Using this regression, for each brain region in each CFC mouse, we calculated a predicted TRAP count of a non-shocked age-matched brain based on the total number of TRAPed cells. From this, we calculated the difference between the observed and predicted value for each region. We then divided this number by the distance from the y axis to account for greater deviations from the predicted value as the distance from the y axis increased. This number was also divided by region volume in voxels to account for variable size of brain regions. This final value, which we term the “learning index”, results in a positive value if a brain region has higher counts than the predicted value of a non-shocked brain and a negative value if the counts are lower than would be predicted.

For voxel-based analysis, the same principle was applied. Cell density per voxel was first calculated as a gaussian distribution of the cell density in a 250um radius surrounding the voxel. In the same way as described above, a learning index was calculated for each voxel to describe whether the observed cell density is higher or lower than what would be predicted in an age-matched non-shocked control. For statistical comparisons between ages as reported in Figure 1E, a one-way ANOVA was performed for each voxel followed by Tukey’s multiple comparisons test, and colored voxels indicate p<0.05 for a particular comparison or set of comparisons. For statistical comparisons between shocked and non-shocked controls, a t-test was performed for each voxel, and colored voxels indicate p<0.05.

For region-based analysis, regions were defined by a collapsed version of the LSFM atlas in which maximum granularity was balanced with the need to account for slight differences in registration which would lead to inaccurate quantification of small brain regions. This atlas was cropped on the anterior and posterior ends to match the amount of tissue visible in our data. Fiber tracts, ventricular systems, cerebellum, and olfactory bulb were excluded from analysis.

### Brain Slice Histology and Immunostaining

Mice were transcardially perfused with phosphate-buffered saline (PBS) followed by 4% paraformaldehyde (PFA) in PBS. Brains were dissected, post-fixed in 4% PFA for 12–24 hours and placed in 30% sucrose for 24–48 hours. They were then embedded in Optimum Cutting Temperature (OCT, Tissue Tek) and stored at −80°C until sectioning. 60 µm floating sections were collected into PBS. Sections were washed 3×10 min in PBS and then blocked in 0.3% PBST containing 10% normal donkey serum (catalog #17-000-121, Jackson Immunoresearch) for 2 hours. Sections were then incubated in a carrier solution (3% donkey serum in 0.3% PBST) containing the primary antibodies for two nights at 4°C. Two days later, sections were washed 3×5 min in PBS and then stained in a carrier solution (5% donkey serum in 0.3% PBST) with secondary antibody for 2 h at room temperature. The following primary antibodies were used: polyclonal chicken anti-GFP (catalog #1020, 1:2000, Aves), monoclonal rabbit anti-RFP (catalog #600-401-379, 1:2000, Rockland), monoclonal rabbit anti-Fos (catalog #226-008, 1:2000, Synaptic Systems), mCherry monoclonal anti-rat (catalog #M11217, 1:500, ThermoFisher), and rabbit polyclonal anti-V5 (Bethyl). The following secondaries were used: Cy2 polyclonal donkey anti-chicken (catalog #703-225-155, 1:1000, Jackson Immunoresearch), Cy3 polyclonal donkey anti-rabbit (catalog #711-005-152, 1:1000, Jackson Immunoresearch), Alexa647 polyclonal donkey anti-rabbit (catalog #711-605-152, 1:1000, Jackson Immunoresearch), and Cy3 polyclonal donkey anti-rat (catalog #712-165-153, 1:1000, Jackson Immunoresearch). Sections were then washed 5 min with PBS, 15 min with PBS+DAPI (catalog #D1306, 1:4000, Thermofisher), and then 5 min with PBS. Sections were mounted on glass slides using FluoroMount-G (catalog #00-4958-02, ThermoFisher) and then imaged at 10x with a Zeiss 700 confocal microscope or at 10x on a Leica DM6 B scanning microscope.

### DREADD Studies

To activate neurons, we used a cre-dependent virus expressing the excitatory DREADD (pAAV-hSyn-DIO-hM3D(Gq)-mCherry) or a fluorophore control (pAAV-hSyn-DIO-mCherry) in TRAP2 mice two weeks prior to fear-conditioning. Viral expression was driven exogenously by intraperitoneally delivered 4-hydroxytamoxifen, which induced fos-dependent expression of cre-recombinase in the TRAP2 mice, allowing specific tagging and long-term labeling of memory encoding ensembles for weeks later. To determine the effects of chronic activation of TRAPed RSC ensembles during development (i.e. comparing freezing levels between retrieval days 1 and 7), 4-hydroxytamoxifen was intraperitoneally injected immediately after 1-day retrieval to tag ensembles involved in retrieval and consolidation (Figure 4) or immediately after contextual fear conditioning (Figure 5).

30 min prior to their first retrieval session, mice were intraperitoneally injected with CNO dissolved in 0.9% saline (1mg/kg, i.p., Fisher Scientific). After the first retrieval session, mice were returned to their home cage. A subset of mice were also tested in an alternate context 3 hours after the first testing session, and then returned to their home cage. Following the first retrieval test, mice were injected daily with CNO (2 mg/kg, i.p.) for 5 days and then returned to the homecage. 30 min prior to the last retrieval day, CNO (1 mg/kg, i.p.) was injected prior to testing in a similar manner as the first retrieval day. Mice were perfused 1 hour after the last retrieval session and their brains collected and processed for Fos immunostaining.

### Fos Analysis in 2-D sections

#### Alignment

Following Fos immunostaining and imaging as described above, sections were aligned to the Allen Brain Atlas^69^ using DeepSlice^70^, a deep-learning based algorithm for automatically calculating the position and slicing angle of each section in relation to the atlas. The DAPI channel was used for alignment. Alignments were imported to the ABBA plugin for FIJI^71,72^ (https://biop.github.io/ijp-imagetoatlas/) which was used to manually check the alignment for each section. ROIs from ABBA were then exported to FIJI, and ROIs for regions to be analyzed were manually checked for tissue damage and image quality.

#### Fos^+^ Cell Detection

Images of the Fos channel were processed using a custom deep learning pipeline. This pipeline has largely the same architecture as our existing 3D pipeline, DeepCOUNT^30^, but uses a 2D U-Net instead of a 3D U-Net. A network for segmenting Fos^+^ cells was trained and validated using manually annotated 100×100 pixel images. Following model training, each section was segmented for Fos^+^ cells. Cells were quantified using 2D maxima detection on the segmented images in FIJI. The number of detected cells in each region was divided by the area of that region to generate Fos density counts.

### MORF+ Cell Detection and Analysis

To quantify dendritic spines of TRAPed cells after learning, 3-5 random ROIs were imaged in the RSP region for each mouse for FIJI analysis. The number of spines were counted along secondary dendrites of TRAPed cells (2 spines per cell, 3-4 cells per mouse) to calculate spine density. Only neurons with their complete morphologies contained within the section were analyzed.

### Quantification and Statistical Analysis

Results were plotted and tested for statistical significance with MATLAB, R and GraphPad Prism v10. Statistical tests are described in the figure legends. Multiple comparisons with single variables were analyzed with one-way ANOVA with post-hoc Tukey test (comparing the mean of each column with that of every other column). For multiple comparisons with more than one variable, two-way ANOVA with post hoc Tukey or Sidak test was used. The relationships between TRAPed cell counts and freezing levels were analyzed using a behavior partial least-squares analysis^33,73,74^ implemented in R. Briefly, a matrix of freezing behavior parameters with each the same number of rows as subjects was multiplied to the regional count matrix (as many lines as subjects and columns as the number of brain regions analyzed) normalized by the total number of labeled cells per region. The product matrix underwent singular value decomposition, resulting in two matrices containing contrasts and saliences respectively. Resampling statistics were used to determine the reliability of brain region saliences. The reliability of brain region saliences was determined using bootstrap ratios in which subjects retained their freezing levels but were resampled with replacement 1000 times. Original saliences were divided by the bootstrap-derived standard deviations to generate bootstrap ratios (saliences). Salience above 3 was considered reliable. For the correlations shown in Figure 2, all possible pairwise correlations between the TRAP signal in the 29 regions were determined by computing Pearson correlation coefficients. Each complete set of correlations was displayed as color-coded correlation matrices using MATLAB. The Fisher Z-Transformation (to transform the sampling distribution of Pearson’s R) was performed in MATLAB. No statistical methods were used to predetermine sample size. Sample sizes were determined based on similar published studies. In all figures, graphs show group means superimposed on individual data points from each animal.

## Supplemental Information Titles & Legends

**Figure S1.**
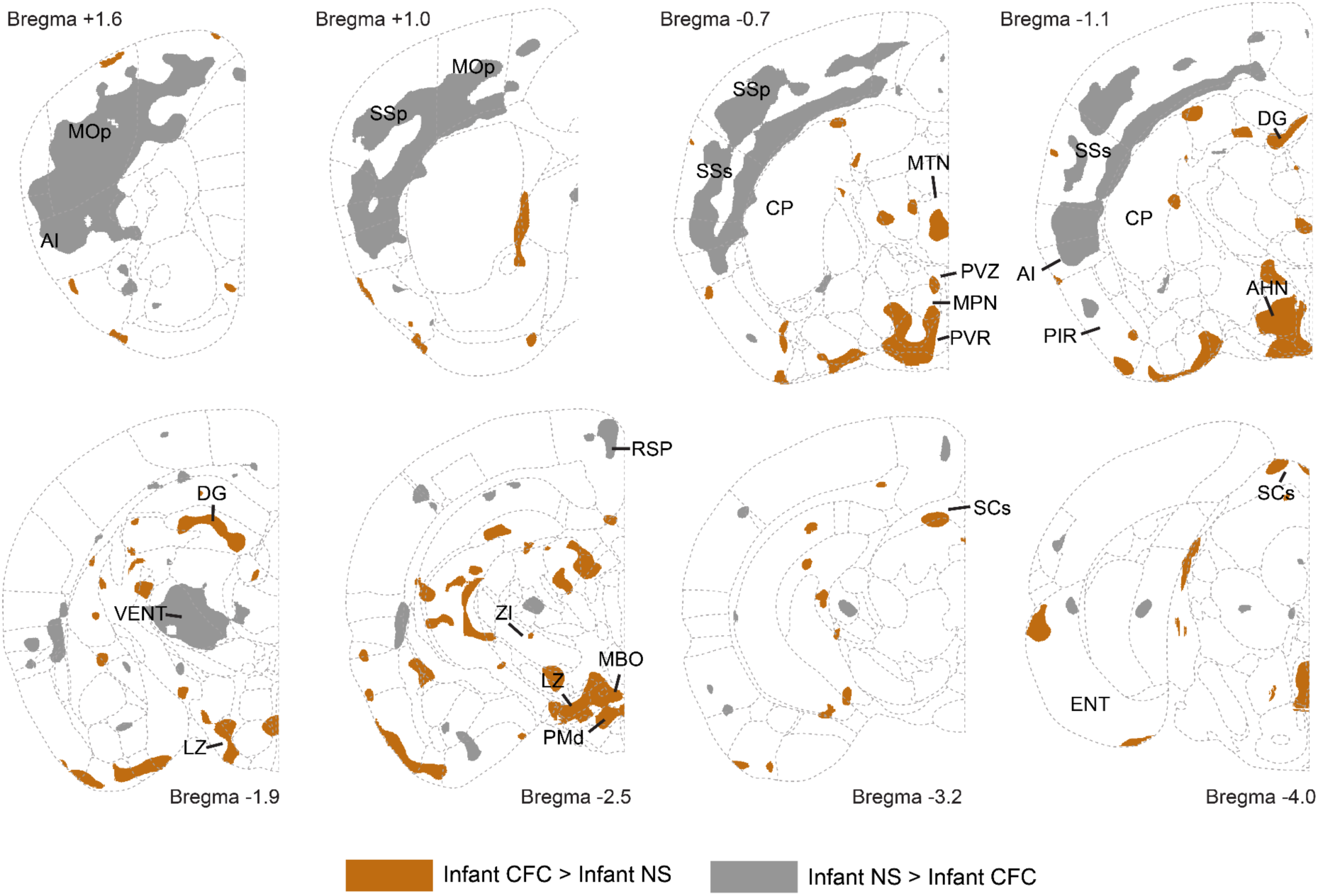
Related to Figure 1. Voxel-based analysis comparing learning indices in infant CFC to age-matched non-shock controls. Values from each voxel underwent an unpaired two-tailed student’s t-test (N=5 CFC, N=4 non-shock). Colored voxels indicate p<0.05 for the comparison indicated in the legend.

**Figure S2.**
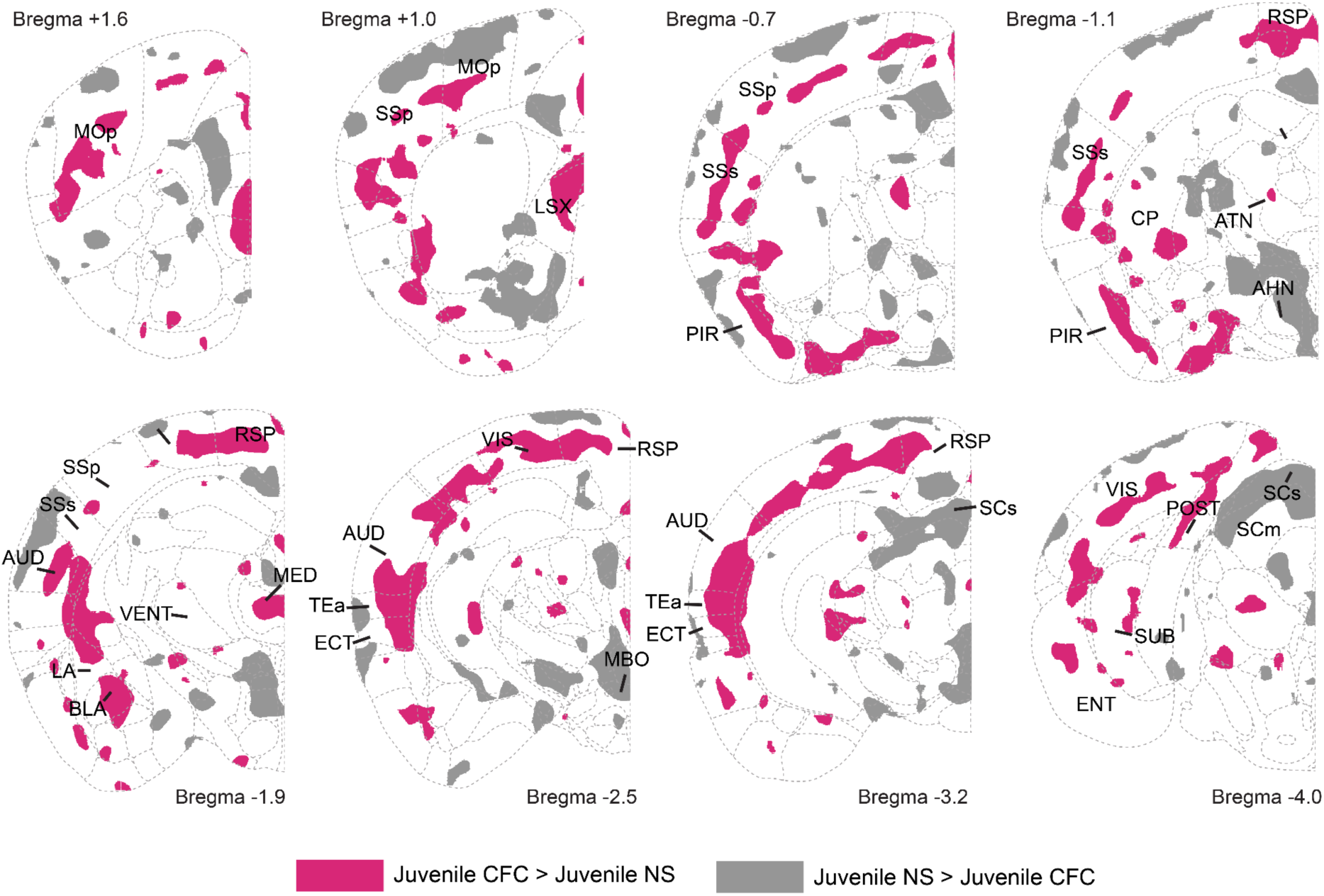
Related to Figure 1. Voxel-based analysis comparing learning indices in juvenile CFC to age-matched non-shock controls. Values from each voxel underwent an unpaired two-tailed student’s t-test (N=4 CFC, N=7 non-shock). Colored voxels indicate p<0.05 for the comparison indicated in the legend.

**Figure S3.**
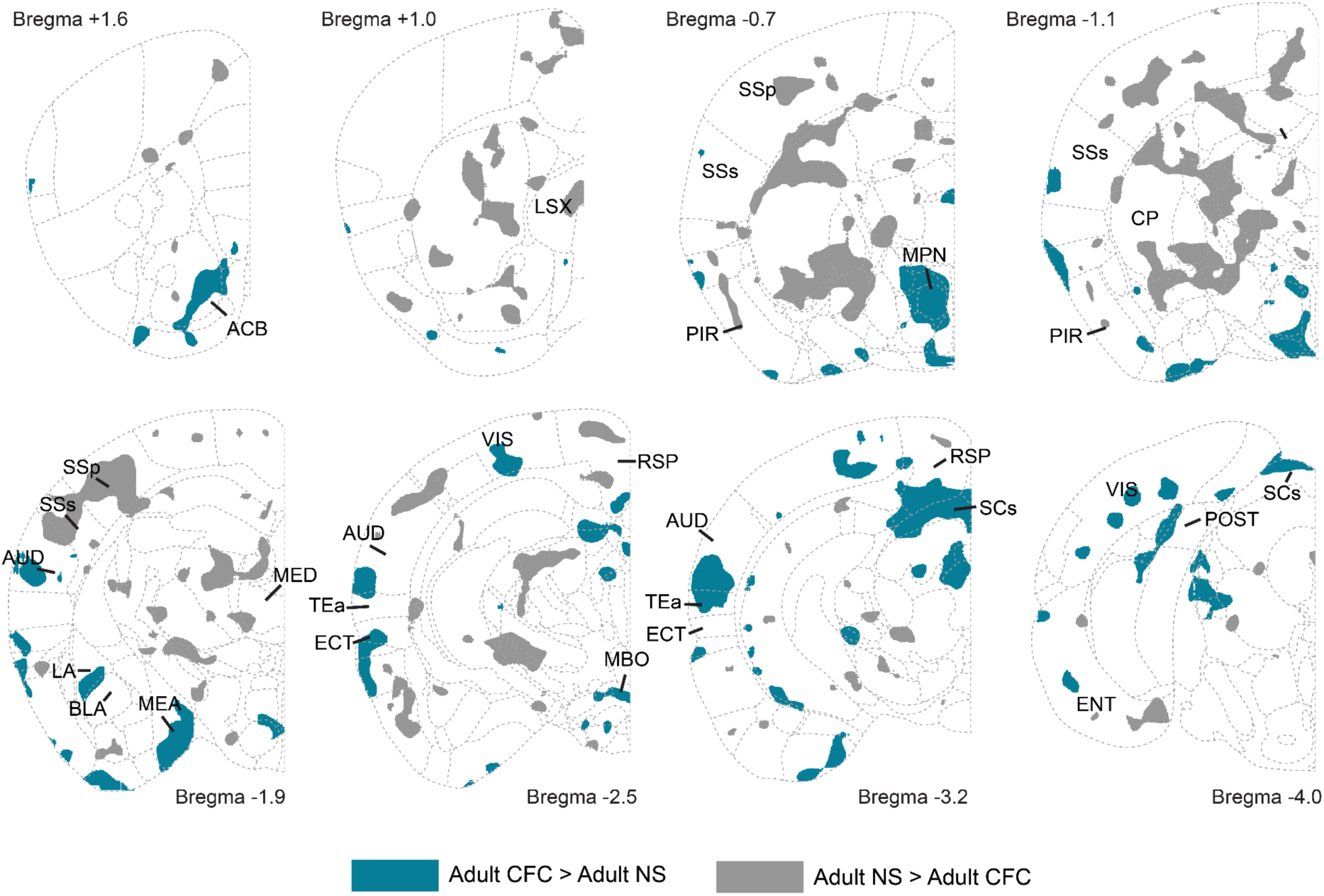
Related to Figure 1. Voxel-based analysis comparing learning indices in adult CFC to age-matched non-shock controls. Values from each voxel underwent an unpaired two-tailed student’s t-test (N=4 CFC, N=7 non-shock). Colored voxels indicate p<0.05 for the comparison indicated in the legend.

**Figure S4.**
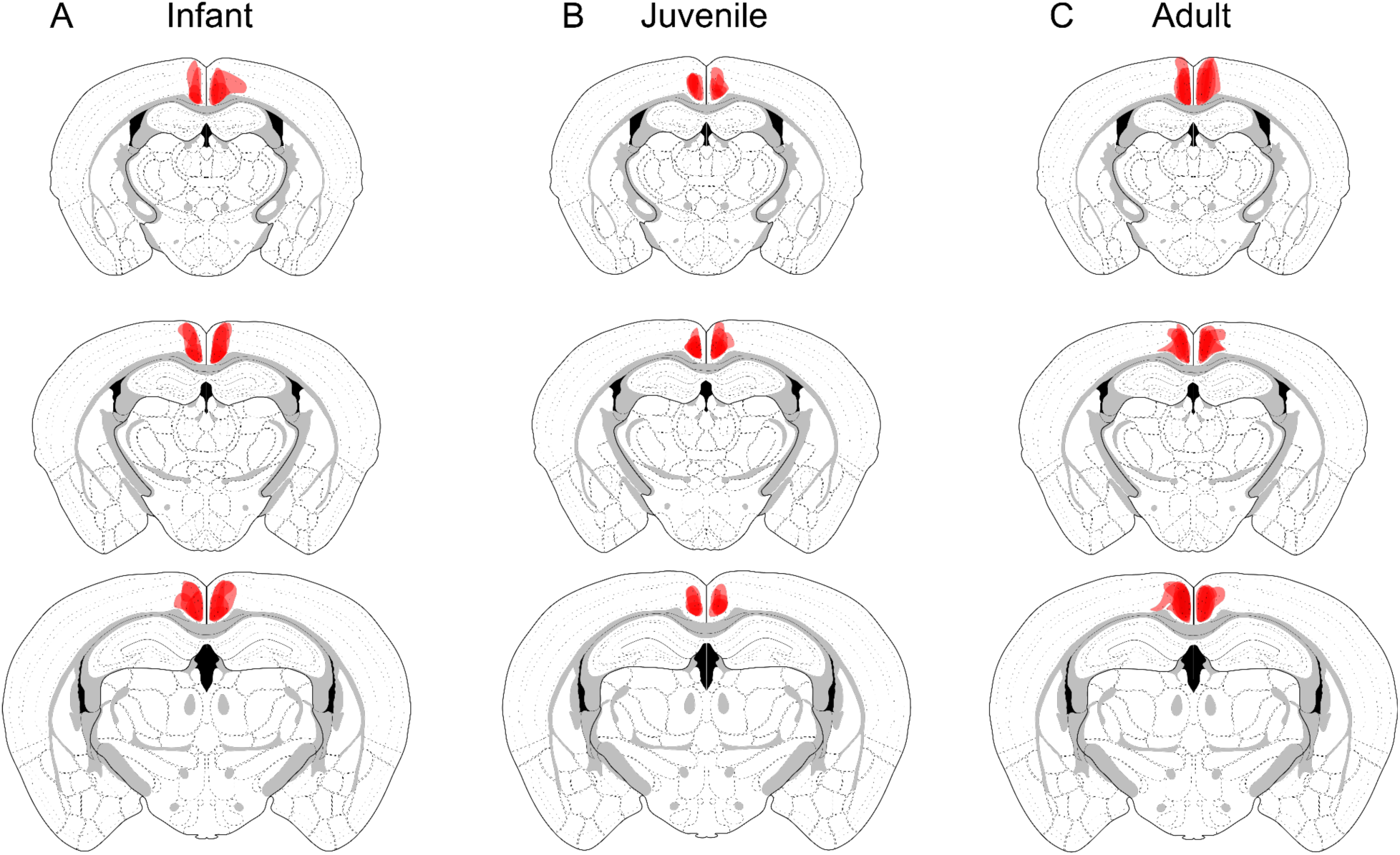
Mapping viral targeting in RSP. Related to Figure 2. (A) Coronal sections showing location of RSP hM3D-mCherry expression in infant-TRAPed mice (B) Coronal sections showing location of RSP hM3D-mCherry expression in juvenile-TRAPed mice (C) Coronal sections showing location of RSP hM3D-mCherry expression in adult-TRAPed mice

**Figure S5.**
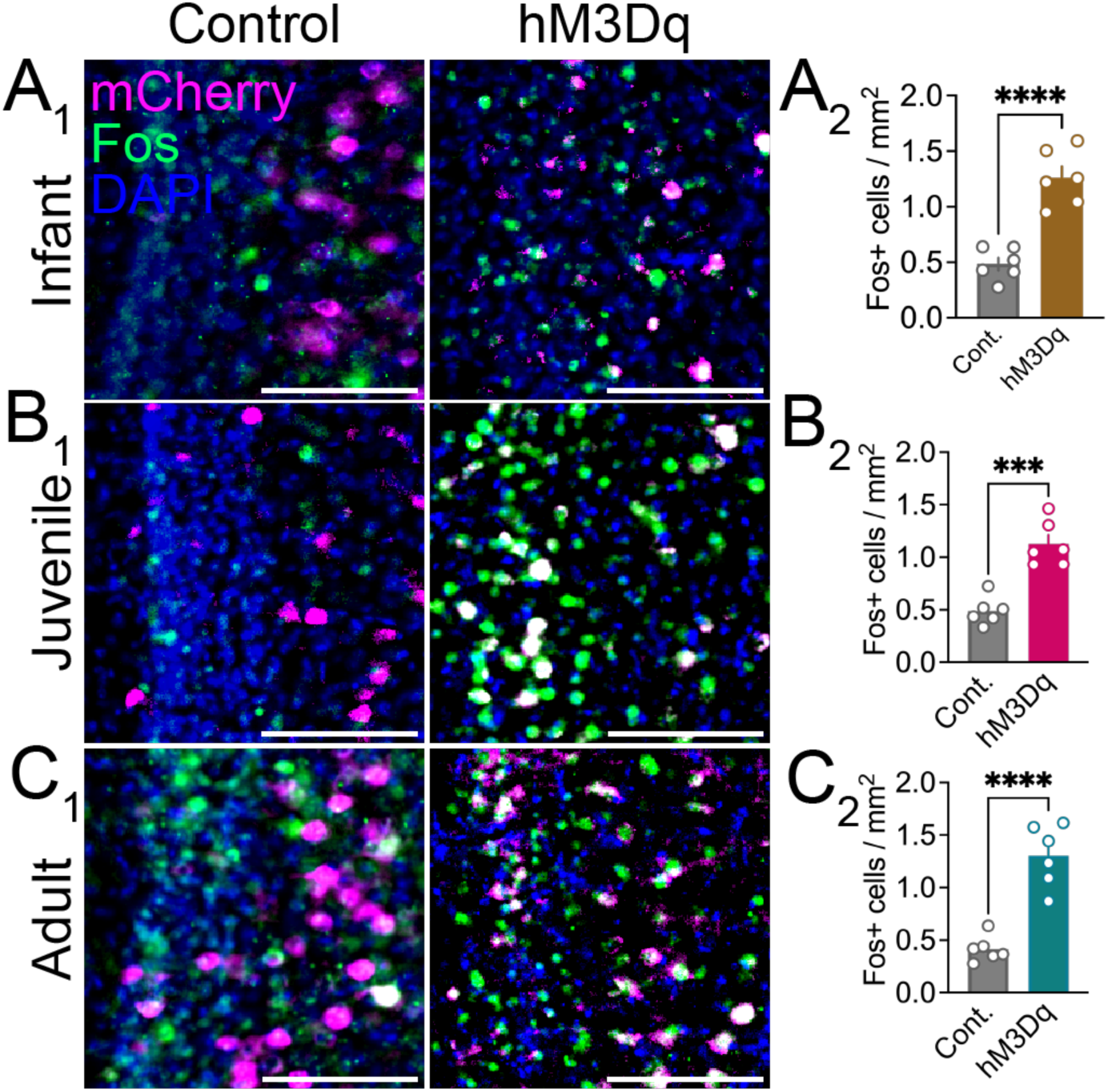
RSP Fos levels in hM3D-expressing vs. control mice. Related to Figure 2. (A_1_) Example images RSP showing AAV-mCherry and Fos+ RSP cells infant-TRAPed mice. (A_2_). Quantification of Fos+ cell density in RSP for infant-TRAPed mice (Cont: N=4, hM3Dq: N=5, P=0.003, two-tailed t-test). (B_1_). Example images RSP showing AAV-mCherry and Fos+ RSP cells infant-TRAPed mice. in juvenile-TRAPed mice (Cont: N=6, hM3Dq: N=6, P=0.0001, two-tailed t-test). (B_2_). Quantification of Fos+ cell density in RSP for juvenile-TRAPed mice. (C_1_). Example images RSP showing AAV-mCherry and Fos+ RSP cells adult-TRAPed mice. (C_2_). Quantification of Fos+ cell density in RSP for adult -TRAPed mice (Cont: N=6, hM3Dq: N=6, P<0.0001, two-tailed t-test). Scale bars, 125 µm. Graphs show mean ± SEM.

**Figure S6.**
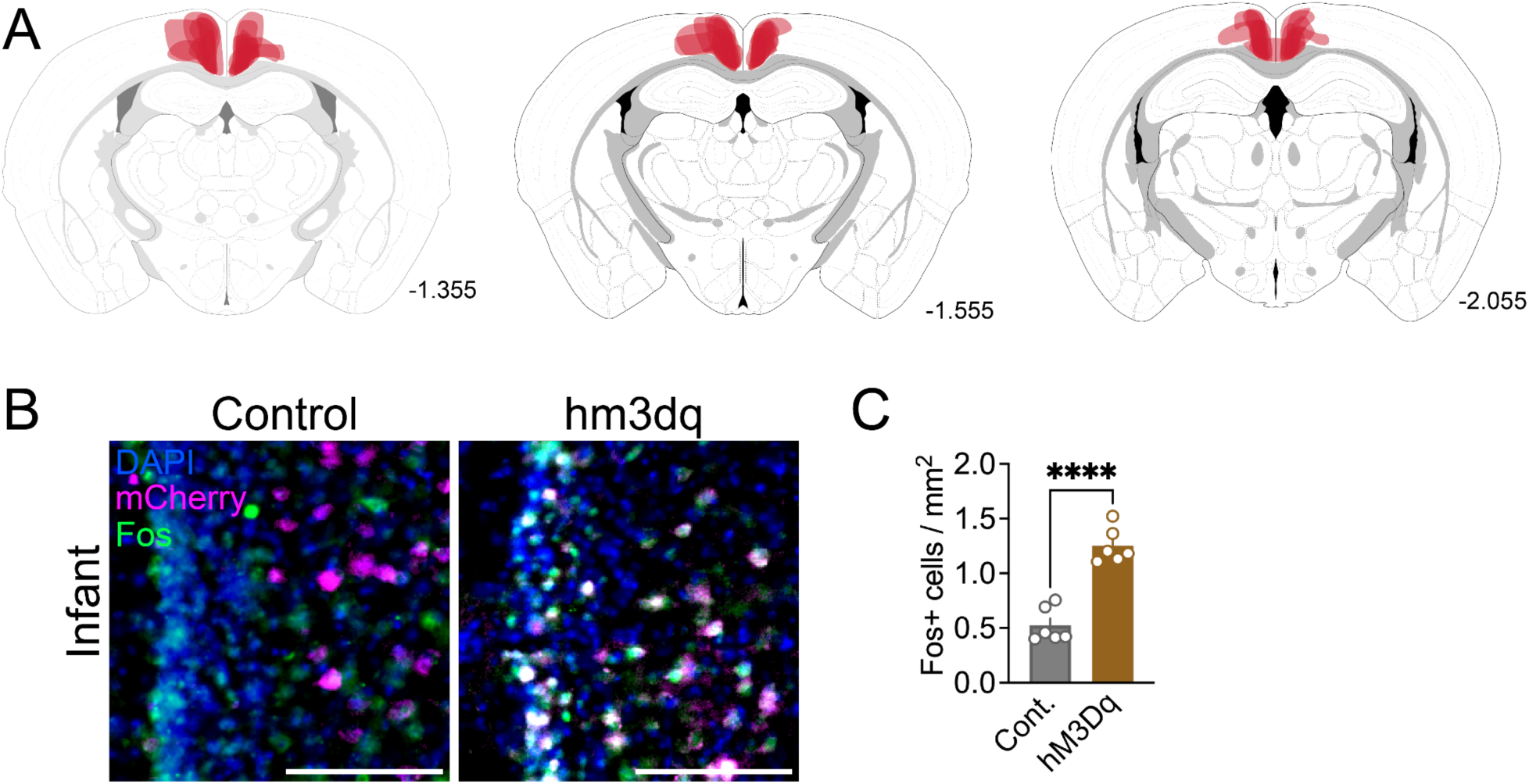
Mapping viral targeting in RSP. Related to Figure 3. (A) Coronal sections showing location of RSP hM3D-mCherry expression in infant-TRAPed mice. (B) Example images RSP showing AAV-mCherry and Fos+ RSP cells infant-TRAPed mice. Scale bars, 125µm. (C) Quantification of Fos+ cell density in RSP for infant-TRAPed mice (Cont: N=6, hM3Dq: N=6, P<0.0001, two-tailed t-test). Graphs show mean ± SEM.

**Figure S7.**
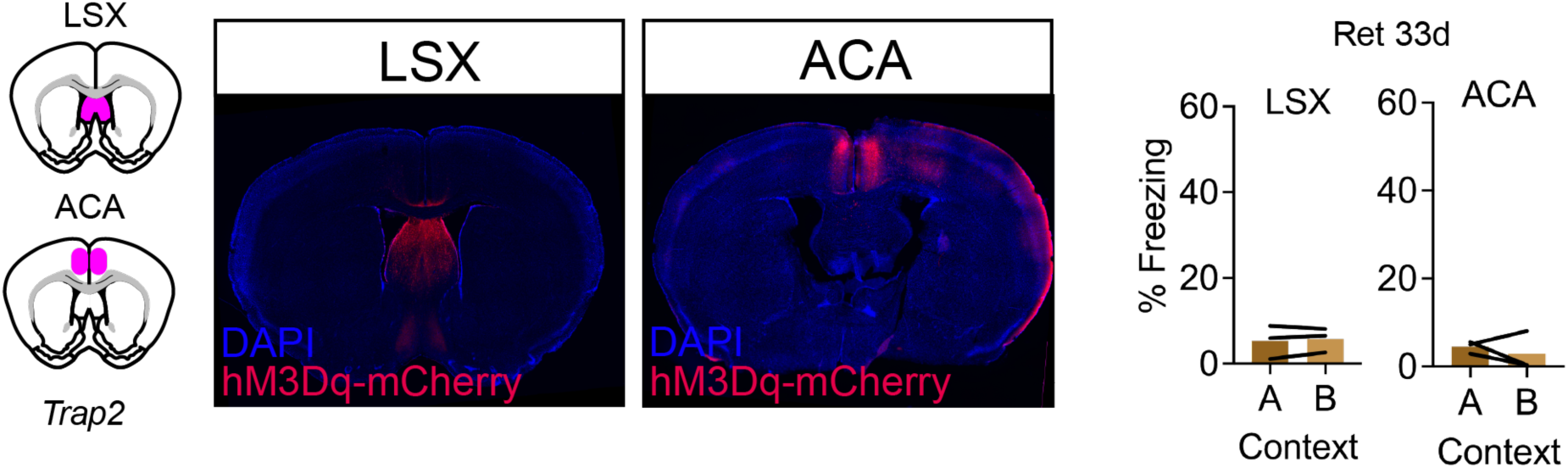
RSP DREADD validation and controls for Figure 3. (A) Representative coronal section showing AAV-hM3Dq injection sites in LSX and ACA. (B) Percent freezing levels in contexts A and B during the 33d retrieval test in LSX- and ACA-targeted mice (LSX: P=0.54, N=3, ACA: P=0.56, N=3, paired t-test). Graphs show mean ± SEM.

## References

1. Bauer, P. J., Tasdemir-Ozdes, A. & Larkina, M. Adults’ reports of their earliest memories: Consistency in events, ages, and narrative characteristics over time. Conscious. Cogn. 27, 76–88 (2014).

2. Campbell, B. A. & Campbell, E. H. Retention and extinction of learned fear in infant and adult rats. J. Comp. Physiol. Psychol. 55, 1–8 (1962).

3. Bauer, P. J. Development of episodic and autobiographical memory: The importance of remembering forgetting. Dev. Rev. DR 38, 146–166 (2015).

4. Lloyd, M. E. & Newcombe, N. S. Implicit memory in childhood: Reassessing developmental invariance. in The development of memory in infancy and childhood, 2nd ed 93–113 (Psychology Press, New York, NY, US, 2009).

5. Ramsaran, A. I., Schlichting, M. L. & Frankland, P. W. The ontogeny of memory persistence and specificity. Dev. Cogn. Neurosci. 36, 100591 (2019).

6. Campbell, B. A. & Spear, N. E. Ontogeny of memory. Psychol. Rev. 79, 215–236 (1972).

7. Callaghan, B. et al. Using a Developmental Ecology Framework to Align Fear Neurobiology Across Species. Annu. Rev. Clin. Psychol. 15, 345–369 (2019).

8. Akers, K. G., Arruda-Carvalho, M., Josselyn, S. A. & Frankland, P. W. Ontogeny of contextual fear memory formation, specificity, and persistence in mice. Learn. Mem. Cold Spring Harb. N 19, 598–604 (2012).

9. Akers, K. G. et al. Hippocampal neurogenesis regulates forgetting during adulthood and infancy. Science 344, 598–602 (2014).

10. Guskjolen, A. Recovery of ‘“Lost”’ Infant Memories in Mice. (2018) doi:10.1016/j.cub.2018.05.059.

11. Guskjolen, A., Josselyn, S. A. & Frankland, P. W. Age-dependent changes in spatial memory retention and flexibility in mice. Neurobiol. Learn. Mem. (2017) doi:10.1016/j.nlm.2016.12.006.

12. Li, S., Kim, J. H. & Richardson, R. Updating memories: Changing the involvement of the prelimbic cortex in the expression of an infant fear memory. Neuroscience 222, 316–325 (2012).

13. Travaglia, A., Bisaz, R., Sweet, E. S., Blitzer, R. D. & Alberini, C. M. Infantile amnesia reflects a developmental critical period for hippocampal learning. Nat. Neurosci. (2016) doi:10.1038/nn.4348.

14. Alberini, C. M. & Travaglia, A. Infantile Amnesia: A Critical Period of Learning to Learn and Remember. J. Neurosci. 37, 5783–5795 (2017).

15. Kim, J. H., Li, S., Hamlin, A. S., McNally, G. P. & Richardson, R. Phosphorylation of mitogen-activated protein kinase in the medial prefrontal cortex and the amygdala following memory retrieval or forgetting in developing rats. Neurobiol. Learn. Mem. 97, 59–68 (2012).

16. Spear, N. E. & Parsons, P. J. Analysis of a Reactivation Treatment: Ontogenetic Determinants of Alleviated Forgetting. in Processes of Animal Memory (PLE: Memory) (Psychology Press, 1976).

17. Campbell, B. A. & Jaynes, J. Reinstatement. 20 (Lawrence Erlbaum Associates Publishers, Mahwah, NJ, US, 2000).

18. Callaghan, B. L. & Richardson, R. Maternal separation results in early emergence of adult-like fear and extinction learning in infant rats. Behav. Neurosci. 125, 20–28 (2011).

19. Bath, K. G., Manzano-Nieves, G. & Goodwill, H. Early life stress accelerates behavioral and neural maturation of the hippocampus in male mice. Horm. Behav. 82, 64–71 (2016).

20. Callaghan, B. L. & Tottenham, N. The Stress Acceleration Hypothesis: Effects of early-life adversity on emotion circuits and behavior. Curr. Opin. Behav. Sci. 7, 76–81 (2016).

21. Frankland, P. W. & Bontempi, B. The organization of recent and remote memories. Nat. Rev. Neurosci. 6, 119–130 (2005).

22. Tonegawa, S., Morrissey, M. D. & Kitamura, T. The role of engram cells in the systems consolidation of memory. Nat. Rev. Neurosci. 19, 485–498 (2018).

23. Frankland, P. W. & Bontempi, B. Fast track to the medial prefrontal cortex. Proc. Natl. Acad. Sci. 103, 509–510 (2006).

24. Klune, C. B., Jin, B. & Denardo, L. A. Linking mPFC circuit maturation to the developmental regulation of emotional memory and cognitive flexibility. eLife 10, (2021).

25. Sugar, J. & Witter, M. P. Postnatal development of retrosplenial projections to the parahippocampal region of the rat. eLife 5, (2016).

26. Haugland, K. G., Sugar, J. & Witter, M. P. Development and topographical organization of projections from the hippocampus and parahippocampus to the retrosplenial cortex. Eur. J. Neurosci. 50, 1799 (2019).

27. DeNardo, L. & Luo, L. Genetic strategies to access activated neurons. Curr. Opin. Neurobiol. 45, 121 (2017).

28. DeNardo, L. A. et al. Temporal evolution of cortical ensembles promoting remote memory retrieval. Nat. Neurosci. 22, 460–469 (2019).

29. Allen, W. E. et al. Thirst-associated preoptic neurons encode an aversive motivational drive. Science 357, 1149–1155 (2017).

30. Gongwer, M. W. et al. Brain-Wide Projections and Differential Encoding of Prefrontal Neuronal Classes Underlying Learned and Innate Threat Avoidance. J. Neurosci. 43, 5810– 5830 (2023).

31. Madisen, L. et al. A robust and high-throughput Cre reporting and characterization system for the whole mouse brain. Nat. Neurosci. 13, 133–140 (2010).

32. Chi, J., Crane, A., Wu, Z. & Cohen, P. Adipo-clear: A tissue clearing method for three-dimensional imaging of adipose tissue. J. Vis. Exp. 2018, 58271 (2018).

33. Wheeler, A. L. et al. Identification of a Functional Connectome for Long-Term Fear Memory in Mice. PLOS Comput. Biol. 9, e1002853 (2013).

34. Kwapis, J. L., Jarome, T. J., Lee, J. L. & Helmstetter, F. J. The retrosplenial cortex is involved in the formation of memory for context and trace fear conditioning. Neurobiol. Learn. Mem. 123, 110–116 (2015).

35. Trask, S., Ferrara, N. C., Jasnow, A. M. & Kwapis, J. L. Contributions of the rodent cingulate-retrosplenial cortical axis to associative learning and memory: A proposed circuit for persistent memory maintenance. Neurosci. Biobehav. Rev. 130, 178–184 (2021).

36. De Sousa, A. F. et al. Optogenetic reactivation of memory ensembles in the retrosplenial cortex induces systems consolidation. Proc. Natl. Acad. Sci. U. S. A. 116, 8576–8581 (2019).

37. Keene, C. S. & Bucci, D. J. Contributions of the retrosplenial and posterior parietal cortices to cue-specific and contextual fear conditioning. Behav. Neurosci. 122, 89–97 (2008).

38. Frankland, P. W., Josselyn, S. A. & Köhler, S. The neurobiological foundation of memory retrieval. Nat. Neurosci. 22, 1576–1585 (2019).

39. Cowansage, K. K. et al. Direct reactivation of a coherent neocortical memory of context. Neuron 84, 432–441 (2014).

40. Armbruster, B. N., Li, X., Pausch, M. H., Herlitze, S. & Roth, B. L. Evolving the lock to fit the key to create a family of G protein-coupled receptors potently activated by an inert ligand. Proc. Natl. Acad. Sci. 104, 5163–5168 (2007).

41. Corcoran, K. A., Frick, B. J., Radulovic, J. & Kay, L. M. Analysis of coherent activity between retrosplenial cortex, hippocampus, thalamus, and anterior cingulate cortex during retrieval of recent and remote context fear memory. Neurobiol. Learn. Mem. 127, 93–101 (2016).

42. Sugar, J., Witter, M., van Strien, N. & Cappaert, N. The Retrosplenial Cortex: Intrinsic Connectivity and Connections with the (Para)Hippocampal Region in the Rat. An Interactive Connectome. Front. Neuroinformatics 5, (2011).

43. van Groen, T. & Michael Wyss, J. Connections of the retrosplenial granular a cortex in the rat. J. Comp. Neurol. 300, 593–606 (1990).

44. van Groen, T. & Wyss, J. M. Connections of the retrosplenial dysgranular cortex in the rat. J. Comp. Neurol. 315, 200–216 (1992).

45. Vann, S. D., Aggleton, J. P. & Maguire, E. A. What does the retrosplenial cortex do? Nat. Rev. Neurosci. 10, 792–802 (2009).

46. Tonegawa, S., Liu, X., Ramirez, S. & Redondo, R. Memory Engram Cells Have Come of Age. Neuron 87, (2015).

47. Josselyn, S. A. & Tonegawa, S. Memory engrams: Recalling the past and imagining the future. Science 367, eaaw4325 (2020).

48. Kitamura, T. et al. Engrams and circuits crucial for systems consolidation of a memory. Science 356, 73–78 (2017).

49. Todd, T. P., Mehlman, M. L., Keene, C. S., DeAngeli, N. E. & Bucci, D. J. Retrosplenial cortex is required for the retrieval of remote memory for auditory cues. Learn. Mem. 23, 278–288 (2016).

50. Baumgärtel, K. et al. PDE4D regulates Spine Plasticity and Memory in the Retrosplenial Cortex. Sci. Rep. 8, 3895 (2018).

51. Frank, A. C. et al. Hotspots of dendritic spine turnover facilitate clustered spine addition and learning and memory. Nat. Commun. 9, 422 (2018).

52. Ramsaran, A. I. et al. A shift in the mechanisms controlling hippocampal engram formation during brain maturation. Science 380, 543–551 (2023).

53. Guskjolen, A. et al. Neurogenesis-mediated circuit remodeling reduces engram reinstatement and promotes forgetting. 2023.10.10.561722 Preprint at 10.1101/2023.10.10.561722 (2023).

54. Zeidler, Z. & DeNardo, L. The Role of Prefrontal Ensembles in Memory Across Time: Time-Dependent Transformations of Prefrontal Memory Ensembles. in Engrams: A Window into the Memory Trace (eds. Gräff, J. & Ramirez, S.) 67–78 (Springer International Publishing, Cham, 2024). doi:10.1007/978-3-031-62983-9_5.

55. Zhou, M. et al. A central amygdala to zona incerta projection is required for acquisition and remote recall of conditioned fear memory. Nat. Neurosci. 21, 1515–1519 (2018).

56. Wang, W. et al. Coordination of escape and spatial navigation circuits orchestrates versatile flight from threats. Neuron 109, 1848–1860.e8 (2021).

57. Miller, A. M. P., Vedder, L. C., Law, L. M. & Smith, D. M. Cues, context, and long-term memory: the role of the retrosplenial cortex in spatial cognition. Front. Hum. Neurosci. 8, (2014).

58. Maviel, T., Durkin, T. P., Menzaghi, F. & Bontempi, B. Sites of Neocortical Reorganization Critical for Remote Spatial Memory. Science 305, 96–99 (2004).

59. Wolbers, T. & Büchel, C. Dissociable Retrosplenial and Hippocampal Contributions to Successful Formation of Survey Representations. J. Neurosci. 25, 3333–3340 (2005).

60. Staresina, B. P., Alink, A., Kriegeskorte, N. & Henson, R. N. Awake reactivation predicts memory in humans. Proc. Natl. Acad. Sci. 110, 21159–21164 (2013).

61. Opalka, A. N. & Wang, D. V. Hippocampal efferents to retrosplenial cortex and lateral septum are required for memory acquisition. doi:10.1101/lm.051797.120.

62. Katche, C. et al. On the role of retrosplenial cortex in long-lasting memory storage. Hippocampus 23, 295–302 (2013).

63. Yasuno, F. et al. Resting-state synchrony between the retrosplenial cortex and anterior medial cortical structures relates to memory complaints in subjective cognitive impairment. Neurobiol. Aging 36, 2145–2152 (2015).

64. Düzel, E., Schütze, H., Yonelinas, A. P. & Heinze, H. Functional phenotyping of successful aging in long-term memory: Preserved performance in the absence of neural compensation. Hippocampus 21, 803–814 (2011).

65. Callaghan, B. L., Li, S. & Richardson, R. The elusive engram: what can infantile amnesia tell us about memory? Trends Neurosci. 37, 47–53 (2014).

66. Mathis, A. et al. DeepLabCut: markerless pose estimation of user-defined body parts with deep learning. Nat. Neurosci. 21, 1281–1289 (2018).

67. Gabriel, C. J. et al. BehaviorDEPOT is a simple, flexible tool for automated behavioral detection based on markerless pose tracking. eLife 11, e74314 (2022).

68. Friedmann, D. et al. Mapping mesoscale axonal projections in the mouse brain using a 3D convolutional network. PNAS 11068–11075 (2020).

69. Wang, Q. et al. The Allen Mouse Brain Common Coordinate Framework: A 3D Reference Atlas. Cell 181, 936–953.e20 (2020).

70. Carey, H. et al. DeepSlice: rapid fully automatic registration of mouse brain imaging to a volumetric atlas. Nat. Commun. 14, 5884 (2023).

71. Chiaruttini, N. et al. An Open-Source Whole Slide Image Registration Workflow at Cellular Precision Using Fiji, QuPath and Elastix. Front. Comput. Sci. 3, (2022).

72. Schindelin, J., et al. Fiji: an open-source platform for biological-image analysis. Nat. Methods 9, 676–682 (2012).

73. Krishnan, A., Williams, L. J., McIntosh, A. R. & Abdi, H. Partial Least Squares (PLS) methods for neuroimaging: A tutorial and review. NeuroImage 56, 455–475 (2011).

74. Franceschini, A. et al. Brain-wide neuron quantification toolkit reveals strong sexual dimorphism in the evolution of fear memory. Cell Rep. 42, 112908 (2023).

