## Supplementary material for "A developmental brain-wide screen identifies retrosplenial cortex as a key player in the emergence of persistent memory": Table 1

**Table 1**. List of Abbreviations

| | |
|---|---|
| ACA | Anterior cingulate cortex |
| ACB | Nucleus accumbens |
| AHN | Anterior hypothalamic Nucleus |
| AI | Agranular Insular Area |
| ATN | Anterior group of the dorsal thalamus |
| AUD | Auditory areas |
| APN | Anterior pretectal nucleus |
| BLA | Basolateral amygdala |
| BMA | Basomedial amygdala |
| BST | Bed nucleus of the stria terminalis |
| CA1 | Hippocampal field CA1 |
| CA2 | Hippocampal field CA2 |
| CA3 | Hippocampal field CA3 |
| CEA | Central amygdalar nucleus |
| CLA | Claustrum |
| CNU | Cerebral nuclei |
| COA | Cortical amygdalar area |
| CP | Caudoputamen |
| CTXsp | Cortical subplate |
| DG | Dentate Gyrus |
| DP | Dorsal peduncular area |
| ECT | Ectorhinal cortex |
| ENTl | Entorhinal area, lateral part |
| ENTm | Entorhinal area, medial part |
| EP | Endopiriform nucleus |
| EPI | Epithalamus |
| FRP | Frontal pole, cerebral cortex |
| GENd | Geniculate group, dorsal thalamus |
| GENv | Geniculate group, ventral thalamus |
| GPe | Globus pallidus, external segment |
| GPi | Globus pallidus, internal segment |
| GU | Gustatory areas |
| HB | Hindbrain |
| HPF | Hippocampal formation |
| IC | Inferior colliculus |
| IGL | Intergeniculat leaflet of the lateral geniculate complex |
| IL | Infralimbic cortex |
| ILM | Intralaminar nucleus of the dorsal thalamus |
| Isoctx | Isocortex |
| LA | Lateral amygdalar nucleus |
| LAT | Lateral group of the dorsal thalamus |
| LSX | Lateral septal complex |
| LZ | Hypothalamic lateral zone |
| MA | Magnocellular Nucleus |
| MB | Midbrain |
| MBO | Mammillary body |
| MBmot | Midbrain, motor related |
| MBsen | Midbrain, sensory related |
| MEA | Medial amygdala |
| MED | Medial group of the dorsal thalamus |
| MOp | Primary motor area |
| MOs | Secondary motor area |
| mPFC | medial prefrontal cortex |
| MPN | Medial preoptic nucleus |
| MPT | Medial pretectal area |
| MS | Medial septal nucleus |
| MTN | Midline group of the dorsal thalamus |
| NDB | Nucleus of the diagonal band |
| NLOT | Nucleus of the lateral olfactory tract |
| OLF | Olfactory areas |
| ORB | Orbital area |
| OT | Olfactory tubercle |
| PA | Posterior amygdalar nucleus |
| PAA | Piriform-amygdalar area |
| PAG | Periaqueductal grey |
| PAL | Pallidum |
| PAR | Parasubiculum |
| PERI | Perirhinal area |
| PH | Posterior hypothalamic nucleus |
| PIR | Piriform area |
| PL | Prelimbic area |
| PMd | Dorsal premammilary nucleus |
| PMv | Ventral premammilary nucleus |
| POST | Postsubiculum |
| PP | Peripeduncular nucleus |
| PPN | Pedunculopontine nucleus |
| PRC | Precommisural nucleus |
| PRE | Presubiculum |
| ProS | Prosubiculum |
| PTLp | Posterior parietal association areas |
| PVHd | Paraventricular hypothalamic nucleus, descending division |
| PVR | Periventricular region |
| PVZ | Periventricular zone |
| RAmb | Midbrain raphe nuclei |
| RSP | Retrosplenial cortex |
| RT | Reticular nucleus of the thalamus |
| sAMY | Striatum-like amygdala |
| SCm | Superior colliculus, motor related |
| SCs | Superior colliculus, sensory related |
| SI | Substantia innominata |
| SNc | Substantia nigra, compact part |
| SNr | Substantia nigra, reticular part |
| SPA | Subparafascicular area |
| SPF | Subparafascicular nucleus |
| SSp | Primary somatosensory area |
| SSs | Secondary somatosensory area |
| STRv | Striatum ventral region |
| SUB | Subiculum |
| TEa | Temporal association area |
| TH | Thalamus |
| TR | Postpiriform transition area |
| TT | Taenia Tecta |
| VENT | Ventral group of the dorsal thalamus |
| VIS | Visual areas |
| VMH | Ventromedial hypothalamic nucleus |
| VTA | Ventral tegmental area |
| ZI | Zona incerta |
