## Supplementary material for "A developmental brain-wide screen identifies retrosplenial cortex as a key player in the emergence of persistent memory": Table 2

Table 2. Detailed statistics for whole-brain map P17 mice.

| Learning Indices | P17 CFC | P17 CFC | P17 CFC | P17 CFC | P17 CFC | P17 NS | P17 NS | P17 NS | P17 NS | Difference CFC-NS | p value (T-test) | Salience |
|---|---|---|---|---|---|---|---|---|---|---|---|---|
| **FRP** | -1.69E-09 | -1.95E-09 | -1.52E-09 | -2.37E-09 | -2.07E-09 | -8.85E-10 | 8.60E-10 | 1.36E-11 | 1.58E-11 | -1.92E-09 | 0.001008 | -5.0117 |
| **MOp** | -2.55E-09 | -1.23E-09 | -1.86E-09 | -8.06E-10 | -1.64E-09 | 1.87E-10 | -3.00E-10 | 7.43E-11 | 5.45E-11 | -1.62E-09 | 0.002307 | -8.11461 |
| **MOs** | -1.01E-09 | -5.03E-10 | -6.26E-10 | -4.21E-10 | -8.73E-10 | -4.63E-10 | 1.81E-10 | 1.82E-10 | 1.40E-10 | -6.96E-10 | 0.007498 | -11.0589 |
| **SSp** | -2.87E-09 | -5.27E-10 | -6.40E-10 | 9.40E-11 | -4.83E-10 | -1.07E-10 | 2.65E-10 | -1.03E-10 | -7.67E-11 | -8.80E-10 | 0.177367 | -1.78918 |
| **SSs** | -7.38E-10 | -3.73E-11 | 2.63E-12 | 3.88E-11 | -5.21E-10 | -3.17E-10 | 2.37E-10 | 5.16E-11 | 4.07E-11 | -2.54E-10 | 0.259206 | -1.56243 |
| **GU** | -2.48E-09 | -2.18E-09 | -2.33E-09 | -7.77E-10 | -9.18E-10 | -2.25E-10 | 6.87E-10 | -3.02E-10 | -2.25E-10 | -1.72E-09 | 0.007597 | -5.38016 |
| **VISC** | -1.59E-09 | -1.46E-09 | -1.13E-09 | -2.37E-10 | -4.36E-10 | -6.50E-10 | 7.23E-10 | -4.94E-11 | -3.29E-11 | -9.68E-10 | 0.043893 | -3.75219 |
| **AUD** | -1.29E-09 | -1.24E-09 | 1.31E-09 | 5.95E-10 | -8.31E-11 | 1.16E-09 | -1.18E-09 | 1.70E-11 | 5.32E-12 | -1.42E-10 | 0.847815 | -0.23625 |
| **VIS** | -3.79E-09 | 1.52E-10 | 1.83E-09 | -2.08E-09 | 6.82E-10 | 1.07E-09 | 2.81E-10 | -8.77E-10 | -6.65E-10 | -5.94E-10 | 0.639267 | -0.95713 |
| **ACA** | -5.58E-10 | -1.24E-10 | 2.17E-09 | -1.44E-09 | -6.76E-10 | 2.27E-10 | 3.84E-10 | -3.98E-10 | -3.00E-10 | -1.04E-10 | 0.888498 | -0.50751 |
| **PL** | 9.43E-11 | 2.65E-10 | 6.40E-10 | -8.10E-10 | -1.27E-09 | -3.66E-10 | 7.07E-10 | -2.24E-10 | -1.65E-10 | -2.05E-10 | 0.667458 | -0.77146 |
| **ILA** | 2.41E-09 | 7.72E-10 | 2.57E-09 | 4.49E-10 | 1.92E-10 | 5.22E-10 | -6.09E-10 | 5.87E-11 | 4.07E-11 | 1.28E-09 | 0.073546 | 3.295054 |
| **ORB** | 6.97E-10 | 3.39E-10 | 1.71E-09 | 3.35E-10 | -4.11E-10 | -6.58E-10 | 7.84E-10 | -8.46E-11 | -5.93E-11 | 5.38E-10 | 0.289859 | 2.196077 |
| **AI** | -1.38E-09 | -1.40E-09 | -1.15E-09 | -7.76E-10 | -6.70E-10 | 3.30E-10 | -1.73E-10 | -1.02E-10 | -7.84E-11 | -1.07E-09 | 0.001032 | -6.28535 |
| **RSP** | -1.90E-09 | 1.12E-09 | 7.10E-10 | -4.47E-10 | 3.47E-10 | -1.39E-10 | -1.29E-11 | 9.83E-11 | 7.47E-11 | -4.02E-11 | 0.949112 | -0.11836 |
| **PTLp** | -2.85E-09 | 1.09E-09 | 1.74E-09 | 4.68E-10 | 5.29E-10 | 1.44E-09 | -1.27E-10 | -8.49E-10 | -6.46E-10 | 2.42E-10 | 0.817332 | 0.382807 |
| **TEa** | -7.14E-10 | -7.49E-10 | 1.24E-09 | 7.23E-10 | 1.32E-09 | 2.71E-10 | -4.42E-10 | 1.13E-10 | 8.27E-11 | 3.59E-10 | 0.526636 | 0.806257 |
| **PERI** | 1.78E-09 | 5.90E-10 | 1.18E-09 | 8.72E-10 | 2.03E-09 | 3.79E-11 | -6.91E-10 | 4.26E-10 | 3.19E-10 | 1.27E-09 | 0.01221 | 8.924656 |
| **ECT** | -9.89E-10 | 2.15E-10 | 1.09E-09 | 3.44E-10 | 1.40E-09 | -3.53E-10 | -1.34E-10 | 3.17E-10 | 2.40E-10 | 3.95E-10 | 0.447188 | 1.132257 |
| **TT** | 6.65E-10 | -1.44E-09 | 8.07E-10 | 4.17E-11 | -1.82E-10 | -2.94E-10 | -4.09E-10 | 4.57E-10 | 3.45E-10 | -4.60E-11 | 0.92818 | -0.22734 |
| **DP** | 3.05E-09 | 1.24E-09 | 2.02E-09 | 9.19E-10 | -1.47E-10 | 4.15E-11 | -1.60E-10 | 7.71E-11 | 5.76E-11 | 1.41E-09 | 0.053777 | 3.514499 |
| **PIR** | 9.04E-10 | 1.85E-11 | 1.56E-09 | 8.84E-10 | 1.35E-09 | -5.99E-10 | 3.47E-10 | 1.63E-10 | 1.26E-10 | 9.34E-10 | 0.032722 | 4.254909 |
| **NLOT** | 1.24E-09 | -1.11E-09 | -4.93E-11 | 1.31E-09 | -1.96E-10 | -1.64E-09 | 8.27E-10 | 5.24E-10 | 4.04E-10 | 2.10E-10 | 0.778886 | 0.696312 |
| **COA** | 3.33E-10 | -7.02E-11 | 1.49E-09 | 2.22E-09 | 2.09E-09 | -1.73E-10 | 1.55E-10 | 1.12E-11 | 9.51E-12 | 1.21E-09 | 0.055374 | 2.337439 |
| **PAA** | 2.35E-09 | 5.75E-10 | 1.16E-09 | 1.49E-09 | 1.90E-09 | -1.05E-09 | 7.41E-10 | 2.01E-10 | 1.57E-10 | 1.48E-09 | 0.017493 | 5.396029 |
| **TR** | 2.38E-09 | 8.41E-10 | 1.91E-09 | 2.40E-09 | 2.46E-09 | -4.14E-10 | -2.70E-11 | 2.86E-10 | 2.17E-10 | 1.98E-09 | 0.001108 | 5.078681 |
| **CA1** | 1.63E-09 | 5.75E-10 | 5.11E-10 | 1.01E-11 | 1.78E-10 | 9.56E-11 | -8.26E-11 | -8.20E-12 | -6.76E-12 | 5.81E-10 | 0.113678 | 2.245678 |
| **CA2** | 2.96E-09 | 6.40E-10 | 3.56E-10 | -4.58E-11 | 2.57E-10 | 1.67E-10 | -3.30E-10 | 1.07E-10 | 7.90E-11 | 8.27E-10 | 0.225711 | 1.648691 |
| **CA3** | 3.59E-09 | 1.17E-09 | 1.11E-09 | 2.02E-10 | 4.02E-10 | -5.52E-11 | -1.66E-10 | 1.44E-10 | 1.08E-10 | 1.29E-09 | 0.1037 | 2.431627 |
| **DG** | 2.42E-09 | 5.32E-10 | 3.24E-11 | 4.06E-10 | 4.68E-10 | -8.14E-11 | -2.01E-10 | 1.84E-10 | 1.38E-10 | 7.63E-10 | 0.159858 | 2.165992 |
| **ENTl** | 4.52E-10 | 5.46E-11 | 1.56E-09 | -1.75E-10 | 2.96E-10 | -1.22E-10 | -1.36E-10 | 1.68E-10 | 1.26E-10 | 4.29E-10 | 0.25759 | 1.40654 |
| **ENTm** | 8.06E-10 | -4.14E-10 | 1.14E-09 | 3.80E-10 | 1.36E-09 | 1.59E-10 | -6.48E-10 | 3.19E-10 | 2.38E-10 | 6.36E-10 | 0.161614 | 2.377178 |
| **PAR** | 1.19E-10 | -9.63E-10 | -2.57E-11 | 3.21E-10 | 9.77E-10 | -2.34E-10 | -5.14E-10 | 4.87E-10 | 3.67E-10 | 5.94E-11 | 0.88957 | 0.10999 |
| **POST** | -4.28E-10 | 5.45E-09 | 2.13E-09 | 3.41E-09 | 2.65E-09 | 7.94E-10 | -6.38E-10 | -9.94E-11 | -7.96E-11 | 2.65E-09 | 0.048775 | 2.655348 |
| **PRE** | 6.24E-10 | 4.66E-10 | -6.27E-10 | 1.44E-09 | 1.49E-09 | 4.08E-10 | -8.28E-10 | 2.75E-10 | 2.04E-10 | 6.65E-10 | 0.229493 | 1.90718 |
| **SUB** | 2.14E-09 | 2.29E-09 | 6.46E-10 | 3.41E-10 | 8.14E-11 | 1.89E-10 | -2.41E-10 | 3.44E-11 | 2.46E-11 | 1.10E-09 | 0.078278 | 2.56867 |
| **ProS** | 1.91E-09 | 4.26E-10 | 3.64E-10 | -2.15E-10 | -1.06E-10 | 9.33E-11 | -6.29E-11 | -1.95E-11 | -1.52E-11 | 4.78E-10 | 0.305814 | 1.275454 |
| **CLA** | 4.64E-10 | 5.34E-10 | 7.66E-10 | 1.06E-09 | 1.05E-09 | -9.99E-10 | 4.35E-10 | 3.65E-10 | 2.80E-10 | 7.55E-10 | 0.056851 | 4.179172 |
| **EP** | 2.53E-09 | -7.66E-11 | 9.89E-10 | 3.39E-10 | -1.91E-11 | -2.55E-10 | -5.31E-11 | 2.00E-10 | 1.51E-10 | 7.42E-10 | 0.223675 | 1.849473 |
| **LA** | 1.27E-09 | 2.38E-10 | 1.42E-09 | -5.38E-10 | -2.38E-10 | -4.72E-10 | -1.58E-10 | 4.09E-10 | 3.10E-10 | 4.09E-10 | 0.424933 | 0.914953 |
| **BLA** | 8.30E-10 | -1.07E-09 | 1.68E-10 | -3.11E-10 | -5.20E-10 | -2.63E-10 | 6.71E-10 | -2.67E-10 | -1.99E-10 | -1.66E-10 | 0.701501 | -0.80912 |
| **BMA** | -1.27E-09 | -1.30E-09 | 2.49E-10 | 9.18E-10 | -3.06E-10 | -2.10E-10 | 1.06E-10 | 6.69E-11 | 5.15E-11 | -3.45E-10 | 0.506065 | -0.49637 |
| **PA** | 1.52E-09 | -1.12E-09 | 8.90E-10 | 1.93E-09 | 1.97E-09 | 1.10E-10 | -4.69E-10 | 2.34E-10 | 1.75E-10 | 1.03E-09 | 0.167827 | 1.751671 |
| **CP** | 4.39E-09 | -4.13E-10 | -2.12E-10 | -6.67E-10 | -3.13E-10 | 3.37E-10 | -1.89E-10 | -9.60E-11 | -7.41E-11 | 5.62E-10 | 0.623161 | 0.480328 |
| **STRv** | 2.85E-09 | -7.87E-10 | -3.42E-10 | -5.70E-10 | -1.69E-09 | -6.19E-11 | 4.61E-10 | -2.60E-10 | -1.95E-10 | -9.33E-11 | 0.919434 | -0.2186 |
| **ACB** | 3.24E-09 | -1.45E-10 | 4.48E-10 | -1.64E-10 | -5.39E-10 | -9.00E-11 | 1.03E-10 | -8.61E-12 | -5.88E-12 | 5.68E-10 | 0.489394 | 0.87977 |
| **OT** | 1.29E-09 | -3.73E-10 | 3.48E-10 | 2.74E-10 | 4.82E-10 | -4.34E-11 | 3.51E-10 | -2.01E-10 | -1.50E-10 | 4.15E-10 | 0.2368 | 1.609316 |
| **LSX** | 1.53E-09 | 4.17E-10 | -4.12E-10 | 1.44E-10 | -3.07E-10 | 4.89E-10 | -7.86E-10 | 1.95E-10 | 1.43E-10 | 2.65E-10 | 0.58595 | 1.052159 |
| **sAMY** | 2.31E-09 | -2.09E-09 | -5.43E-10 | 1.25E-09 | -9.53E-10 | -3.70E-10 | 1.91E-10 | 1.16E-10 | 8.92E-11 | -1.15E-11 | 0.990222 | 0.143669 |
| **CEA** | -5.83E-10 | -2.07E-09 | -2.04E-09 | -1.18E-09 | -1.81E-09 | -9.13E-10 | 6.29E-10 | 1.82E-10 | 1.42E-10 | -1.55E-09 | 0.009123 | -5.32125 |
| **MEA** | -3.02E-09 | -2.10E-09 | -1.09E-09 | 2.21E-09 | 7.61E-10 | 5.46E-10 | -5.01E-10 | -2.71E-11 | -2.38E-11 | -6.46E-10 | 0.573975 | -0.39707 |
| **PAL** | 7.77E-09 | -3.44E-10 | 1.15E-09 | 1.86E-10 | 1.02E-09 | -1.43E-10 | -5.67E-11 | 1.30E-10 | 9.84E-11 | 1.95E-09 | 0.282978 | 1.390092 |
| **GPe** | 3.70E-09 | 1.32E-09 | 7.05E-10 | 3.64E-10 | 1.10E-09 | -2.18E-10 | -3.60E-10 | 3.76E-10 | 2.83E-10 | 1.42E-09 | 0.077427 | 2.761713 |
| **GPi** | 4.90E-09 | -3.55E-10 | 3.17E-11 | -2.68E-11 | 1.04E-09 | -4.88E-11 | -3.04E-10 | 2.30E-10 | 1.73E-10 | 1.11E-09 | 0.352064 | 1.140879 |
| **SI** | 7.04E-09 | -2.42E-10 | 7.26E-10 | 5.01E-11 | -4.39E-10 | 8.70E-11 | -1.38E-10 | 3.34E-11 | 2.45E-11 | 1.42E-09 | 0.404699 | 1.099638 |
| **MA** | 1.15E-08 | 1.28E-10 | 9.54E-10 | 6.70E-10 | -8.12E-10 | 2.72E-10 | -2.37E-10 | -2.21E-11 | -1.83E-11 | 2.48E-09 | 0.366541 | 1.268891 |
| **MS** | 4.04E-09 | -2.02E-09 | -1.56E-10 | 1.50E-09 | -2.22E-09 | 1.88E-09 | -5.19E-10 | -8.80E-10 | -6.72E-10 | 2.76E-10 | 0.85371 | 0.404711 |
| **NDB** | 1.12E-08 | -1.51E-09 | 1.08E-09 | 1.80E-10 | -2.12E-11 | 7.12E-10 | -3.42E-10 | -2.38E-10 | -1.83E-10 | 2.21E-09 | 0.426761 | 0.994976 |

| | | | | | | | | | | | | |
|---|---|---|---|---|---|---|---|---|---|---|---|---|
| **TRS** | 4.37E-10 | -1.95E-09 | -1.92E-09 | -4.60E-10 | 3.52E-10 | 9.34E-10 | 1.85E-10 | -7.26E-10 | -5.51E-10 | -6.68E-10 | 0.359858 | -1.70832 |
| **BST** | 1.12E-09 | -4.67E-10 | -3.31E-10 | 4.40E-12 | 7.00E-10 | -7.06E-10 | -1.20E-10 | 5.36E-10 | 4.07E-10 | 1.77E-10 | 0.691119 | 0.52808 |
| **VENT** | -4.60E-10 | 3.67E-11 | -5.29E-10 | 4.78E-10 | -1.15E-10 | -4.79E-11 | -4.11E-10 | 2.99E-10 | 2.25E-10 | -1.34E-10 | 0.60926 | -0.14757 |
| **SPF** | -9.95E-09 | -3.72E-09 | -8.90E-09 | 2.22E-09 | -4.74E-10 | -9.21E-10 | 8.75E-10 | 2.75E-11 | 2.65E-11 | -4.17E-09 | 0.164604 | -1.68141 |
| **SPA** | 1.01E-09 | 1.99E-09 | -2.29E-09 | 3.60E-09 | 9.22E-10 | 6.56E-10 | -1.99E-09 | 8.73E-10 | 6.51E-10 | 1.00E-09 | 0.447849 | 1.334777 |
| **PP** | 7.07E-09 | 5.67E-09 | -1.98E-09 | 2.60E-09 | 1.33E-09 | 4.29E-10 | -5.82E-10 | 1.01E-10 | 7.30E-11 | 2.93E-09 | 0.152502 | 2.183945 |
| **GENd** | 2.00E-09 | -7.82E-10 | -1.59E-10 | 6.81E-10 | 2.74E-10 | 7.48E-11 | -6.56E-10 | 3.79E-10 | 2.84E-10 | 3.83E-10 | 0.522473 | 0.969221 |
| **LAT** | 2.65E-10 | 8.21E-11 | 5.50E-11 | -1.31E-10 | -1.06E-10 | -2.96E-10 | -3.23E-11 | 2.13E-10 | 1.62E-10 | 2.14E-11 | 0.873615 | 0.288312 |
| **ATN** | 1.19E-09 | 1.21E-09 | 1.49E-09 | 5.03E-10 | 9.41E-10 | -3.31E-10 | -3.13E-10 | 4.19E-10 | 3.16E-10 | 1.04E-09 | 0.004853 | 7.810505 |
| **MED** | -9.27E-10 | -3.97E-10 | 5.21E-10 | -5.15E-10 | -2.53E-10 | 6.32E-10 | -3.96E-10 | -1.52E-10 | -1.18E-10 | -3.06E-10 | 0.388047 | -1.6703 |
| **MTN** | -1.79E-09 | 1.93E-09 | -2.88E-10 | -1.85E-09 | 1.47E-09 | 1.00E-10 | 1.25E-10 | -1.47E-10 | -1.11E-10 | -9.82E-11 | 0.916193 | -0.39659 |
| **ILM** | -2.28E-09 | -3.71E-10 | 8.84E-10 | -1.18E-09 | 1.75E-10 | -1.62E-09 | 6.04E-10 | 6.60E-10 | 5.06E-10 | -5.90E-10 | 0.478585 | -1.37019 |
| **RT** | 4.71E-09 | 1.73E-11 | 6.69E-10 | 1.31E-10 | 3.93E-10 | 6.74E-11 | -1.09E-10 | 2.76E-11 | 2.03E-11 | 1.18E-09 | 0.279114 | 1.436883 |
| **GENv** | -9.31E-09 | 7.28E-10 | -8.15E-09 | -5.93E-10 | 1.77E-09 | 1.21E-10 | 5.59E-10 | -4.43E-10 | -3.33E-10 | -3.09E-09 | 0.282381 | -1.59887 |
| **EPI** | 5.97E-09 | 2.57E-09 | 7.56E-10 | 1.10E-09 | 1.35E-09 | -1.35E-10 | -3.82E-10 | 3.37E-10 | 2.54E-10 | 2.33E-09 | 0.070387 | 2.877846 |
| **PVZ** | -4.03E-09 | -3.34E-10 | -1.69E-09 | -2.51E-09 | 3.70E-09 | 8.78E-10 | -1.67E-09 | 5.17E-10 | 3.82E-10 | -1.00E-09 | 0.544988 | -1.30766 |
| **PVR** | -3.00E-09 | 2.89E-09 | -9.28E-10 | -1.77E-09 | 1.08E-09 | -9.47E-10 | 1.91E-10 | 4.90E-10 | 3.74E-10 | -3.74E-10 | 0.768516 | -0.45449 |
| **AHN** | 5.75E-09 | 3.58E-09 | 4.20E-09 | -4.37E-10 | 1.60E-09 | 6.03E-10 | -5.07E-10 | -6.08E-11 | -4.95E-11 | 2.94E-09 | 0.048609 | 2.69713 |
| **MBO** | -1.25E-09 | -4.07E-10 | 9.79E-10 | -8.68E-10 | -9.79E-10 | 8.49E-10 | -9.41E-10 | 6.27E-11 | 4.16E-11 | -5.07E-10 | 0.388279 | -1.59474 |
| **MPN** | 9.18E-10 | 4.17E-09 | -5.89E-10 | -1.31E-10 | -2.80E-10 | 1.04E-09 | -9.62E-10 | -5.03E-11 | -4.44E-11 | 8.20E-10 | 0.462706 | 1.081269 |
| **PMd** | 2.43E-09 | 2.37E-09 | 3.64E-09 | 6.34E-09 | 7.79E-09 | -2.14E-09 | 1.01E-09 | 7.30E-10 | 5.61E-10 | 4.47E-09 | 0.014865 | 3.705354 |
| **PMv** | -4.27E-09 | -3.11E-09 | -2.36E-09 | -5.00E-09 | -6.42E-09 | 2.29E-09 | -1.72E-09 | -3.70E-10 | -2.92E-10 | -4.21E-09 | 0.006226 | -5.1612 |
| **PVHd** | -8.96E-09 | 4.64E-09 | -5.26E-09 | 5.96E-10 | 3.30E-09 | -1.53E-09 | 1.02E-09 | 3.29E-10 | 2.57E-10 | -1.16E-09 | 0.708969 | -0.38111 |
| **VMH** | 2.86E-09 | 8.49E-11 | 1.94E-12 | -8.51E-10 | 1.62E-10 | -1.27E-09 | -7.85E-10 | 1.34E-09 | 1.01E-09 | 3.79E-10 | 0.690222 | 0.481975 |
| **PH** | -8.13E-09 | 3.39E-09 | -3.74E-09 | 7.83E-10 | -5.62E-10 | 6.23E-10 | -5.82E-10 | -2.45E-11 | -2.24E-11 | -1.65E-09 | 0.489027 | -0.66983 |
| **LZ** | 2.41E-09 | 1.62E-10 | 5.80E-10 | 5.07E-10 | 1.95E-10 | -8.54E-10 | 4.93E-10 | 2.33E-10 | 1.80E-10 | 7.58E-10 | 0.203845 | 2.236378 |
| **ZI** | -1.70E-09 | -1.54E-09 | -2.28E-09 | -9.46E-10 | 5.09E-11 | -1.89E-10 | 1.74E-10 | 8.94E-12 | 7.91E-12 | -1.28E-09 | 0.025389 | -4.17286 |
| **MBsen** | 6.45E-09 | 1.85E-09 | -2.75E-10 | -1.92E-10 | 7.49E-10 | -1.87E-10 | -3.07E-10 | 3.21E-10 | 2.42E-10 | 1.70E-09 | 0.269868 | 1.465841 |
| **SCs** | -3.61E-09 | 3.72E-09 | -2.14E-09 | 4.77E-09 | 3.05E-09 | -1.46E-09 | 2.22E-10 | 8.01E-10 | 6.10E-10 | 1.12E-09 | 0.587246 | 0.989319 |
| **IC** | -2.69E-09 | 3.77E-09 | -3.51E-09 | 2.53E-09 | 3.99E-10 | -1.03E-10 | 3.09E-10 | -1.35E-10 | -1.00E-10 | 1.08E-10 | 0.948487 | 0.457602 |
| **MBmot** | 2.94E-10 | 3.79E-10 | -2.46E-09 | 2.07E-10 | -1.30E-09 | -7.77E-11 | 1.41E-10 | -4.13E-11 | -3.05E-11 | -5.73E-10 | 0.400279 | -0.86916 |
| **SNr** | 4.05E-09 | -2.62E-10 | 5.51E-10 | 5.54E-10 | 5.01E-10 | 2.39E-10 | -4.77E-10 | 1.56E-10 | 1.15E-10 | 1.07E-09 | 0.259404 | 1.609762 |
| **VTA** | 7.80E-09 | -6.61E-10 | -7.26E-10 | -8.00E-10 | -2.15E-09 | 1.04E-09 | -8.68E-11 | -6.17E-10 | -4.70E-10 | 7.26E-10 | 0.735368 | 0.383424 |
| **SCm** | -3.37E-09 | 9.44E-10 | -3.21E-09 | 1.66E-09 | 6.32E-11 | -3.26E-10 | 2.40E-10 | 5.46E-11 | 4.31E-11 | -7.83E-10 | 0.534699 | -0.4073 |
| **PAG** | -3.46E-09 | 4.07E-10 | -4.94E-09 | 6.92E-10 | -1.22E-09 | 3.72E-11 | -7.97E-11 | 2.78E-11 | 2.06E-11 | -1.71E-09 | 0.211852 | -1.43657 |
| **PRC** | -2.03E-09 | 5.87E-09 | -2.27E-09 | 1.21E-09 | 2.81E-10 | -1.67E-09 | 1.47E-09 | 1.23E-10 | 1.03E-10 | 6.04E-10 | 0.741885 | 0.719042 |
| **APN** | -2.60E-09 | 5.89E-10 | -1.49E-09 | 1.58E-09 | 5.14E-10 | 8.84E-11 | 5.03E-11 | -9.01E-11 | -6.82E-11 | -2.76E-10 | 0.759602 | -0.02183 |
| **MPT** | 2.02E-09 | 2.56E-09 | -2.01E-09 | 5.07E-09 | 3.05E-09 | 1.34E-09 | -1.47E-09 | 8.74E-11 | 5.70E-11 | 2.13E-09 | 0.17295 | 1.867295 |
| **SNc** | 7.30E-09 | 7.61E-10 | 2.33E-09 | 5.52E-10 | -4.23E-10 | 1.76E-10 | -2.42E-10 | 4.33E-11 | 3.13E-11 | 2.10E-09 | 0.21926 | 1.828234 |
| **PPN** | 2.95E-10 | 1.45E-09 | -5.21E-10 | 1.34E-09 | -1.33E-10 | -6.03E-10 | 6.46E-10 | -2.96E-11 | -1.84E-11 | 4.87E-10 | 0.361258 | 1.531777 |
| **RAmb** | -2.09E-09 | -7.16E-10 | -3.28E-09 | 3.13E-09 | -4.38E-10 | 6.42E-10 | -2.55E-09 | 1.24E-09 | 9.29E-10 | -7.45E-10 | 0.622905 | -0.14001 |
