## Supplementary material for "A developmental brain-wide screen identifies retrosplenial cortex as a key player in the emergence of persistent memory": Table 3

**Table 3. Detailed statistics for whole-brain map P17 mice.**

| Learning Indices | P25 CFC | P25 CFC | P25 CFC | P25 CFC | P25 NS | P25 NS | P25 NS | P25 NS | P25 NS | P25 NS | P25 NS | Difference CFC-NS | p value (T-test) | Salience |
|---|---|---|---|---|---|---|---|---|---|---|---|---|---|---|
| FRP | 1.20E-09 | 1.27E-09 | 1.24E-09 | 8.27E-10 | -2.00E-10 | 4.86E-11 | 2.07E-11 | 3.24E-10 | -1.20E-10 | -1.75E-11 | -5.66E-11 | 1.14E-09 | 0.000065 | 9.527361 |
| MOp | 1.62E-09 | 1.43E-09 | 1.40E-09 | 1.50E-09 | 1.69E-10 | -5.84E-10 | -6.57E-10 | -1.58E-10 | 5.54E-10 | 2.32E-10 | 3.89E-10 | 1.50E-09 | 0.000238 | 8.971715 |
| MOs | 1.64E-09 | 1.29E-09 | 1.34E-09 | 8.81E-10 | 3.34E-10 | -1.82E-10 | -1.92E-10 | -4.10E-10 | 1.99E-10 | 6.06E-11 | 1.82E-10 | 1.29E-09 | 0.000007 | 5.983922 |
| SSp | 1.49E-09 | 3.64E-10 | 1.16E-09 | 1.62E-09 | 4.90E-10 | 1.79E-10 | 2.52E-10 | -6.52E-10 | -1.96E-10 | -1.52E-10 | 1.11E-10 | 1.16E-09 | 0.006832 | 3.940903 |
| SSs | 1.96E-09 | 1.63E-09 | 1.36E-09 | 2.21E-09 | 6.92E-10 | 5.48E-10 | -4.12E-11 | -4.02E-10 | -3.05E-10 | -4.91E-10 | 3.60E-11 | 1.78E-09 | 0.00014 | 6.766449 |
| GU | 1.70E-09 | 2.39E-09 | 1.88E-09 | 2.83E-10 | 1.78E-10 | -6.51E-10 | -2.92E-10 | 8.74E-11 | -3.83E-10 | 1.68E-10 | 8.61E-10 | 1.57E-09 | 0.000042 | 4.35761 |
| VISC | 1.13E-09 | 9.27E-10 | 2.56E-10 | 3.33E-10 | 3.96E-10 | 1.37E-10 | -4.43E-10 | 3.62E-10 | -4.94E-10 | -3.19E-10 | 3.58E-10 | 6.61E-10 | 0.016922 | 3.020346 |
| AUD | 8.22E-10 | 1.02E-09 | 3.53E-09 | 1.85E-09 | 1.34E-09 | 1.48E-09 | 1.66E-11 | -1.67E-09 | 6.23E-10 | -1.09E-09 | -6.28E-10 | 1.80E-09 | 0.027009 | 3.48676 |
| VIS | 2.85E-10 | -3.87E-09 | -1.57E-09 | -4.99E-10 | -3.82E-10 | -3.62E-10 | -2.96E-10 | 4.30E-11 | 4.91E-10 | -1.29E-10 | 5.71E-10 | -1.40E-09 | 0.026241 | -1.81775 |
| ACA | 2.59E-09 | 1.62E-09 | 2.65E-09 | 3.57E-10 | 5.66E-10 | -6.58E-10 | 2.64E-10 | -6.06E-10 | -5.15E-10 | 6.35E-10 | 3.42E-10 | 1.80E-09 | 0.000161 | 3.432584 |
| PL | 9.11E-10 | 1.78E-09 | 7.64E-10 | 2.97E-10 | 8.58E-10 | 6.07E-10 | 7.99E-10 | -7.63E-10 | -1.12E-09 | -3.42E-10 | 5.53E-11 | 9.23E-10 | 0.049408 | 4.350111 |
| ILA | 1.73E-09 | 4.92E-09 | 3.15E-09 | 2.31E-10 | -4.37E-10 | -1.43E-09 | 1.79E-10 | 5.74E-10 | -7.66E-10 | 1.11E-09 | 7.50E-10 | 2.51E-09 | 0.00893 | 3.488942 |
| ORB | 2.58E-09 | 2.86E-09 | 3.20E-09 | -7.82E-10 | 2.92E-10 | 8.41E-10 | 1.89E-09 | -1.18E-09 | -1.27E-09 | -5.10E-10 | 6.73E-11 | 1.94E-09 | 0.000826 | 2.47074 |
| AI | 5.11E-10 | 8.51E-10 | 5.13E-10 | -1.20E-10 | 4.64E-10 | 1.52E-12 | 5.65E-12 | -1.02E-10 | -5.18E-10 | 1.44E-11 | 1.61E-10 | 4.35E-10 | 0.003027 | 2.680634 |
| RSP | 1.42E-09 | 6.88E-10 | 2.07E-09 | -1.29E-10 | 8.26E-11 | -7.76E-10 | -6.87E-10 | 8.39E-12 | 2.01E-10 | 3.85E-11 | 1.06E-09 | 1.02E-09 | 0.030565 | 2.458158 |
| PTLp | 7.22E-10 | -4.52E-09 | -1.20E-09 | 1.21E-09 | 5.24E-10 | 1.90E-10 | 1.34E-10 | -1.88E-10 | -5.81E-10 | -7.59E-11 | 3.91E-11 | -9.53E-10 | 0.055466 | -0.64927 |
| TEa | 2.51E-09 | 2.12E-09 | 3.92E-09 | 2.92E-09 | 8.77E-10 | 1.20E-09 | 2.40E-10 | -8.72E-10 | -8.18E-11 | -8.47E-10 | -4.43E-10 | 2.86E-09 | 0.000549 | 7.351092 |
| PERI | 2.25E-10 | 3.98E-11 | -1.40E-09 | 3.45E-10 | -8.17E-11 | 3.68E-11 | 7.06E-10 | 1.20E-11 | -9.62E-10 | -1.30E-10 | 4.55E-10 | -2.04E-10 | 0.761294 | -0.52934 |
| ECT | 3.49E-09 | 3.03E-09 | 2.54E-09 | 1.77E-09 | 1.68E-10 | 8.51E-11 | 6.97E-10 | -4.76E-10 | -6.28E-10 | -9.30E-11 | 2.89E-10 | 2.70E-09 | 0.000002 | 5.817898 |
| TT | -6.72E-10 | -7.40E-10 | 9.92E-10 | -1.02E-09 | 3.67E-10 | 2.70E-10 | 2.76E-10 | 5.59E-12 | -8.25E-10 | -2.02E-10 | 1.53E-10 | -3.67E-10 | 0.821864 | -0.94871 |
| DP | 2.38E-09 | 4.93E-09 | 4.78E-09 | -2.93E-10 | 7.11E-10 | -2.11E-10 | 4.46E-10 | -4.82E-10 | -7.61E-10 | 6.08E-10 | -2.39E-10 | 2.94E-09 | 0.000024 | 3.06743 |
| PIR | 1.56E-09 | 4.78E-10 | -1.15E-09 | -4.10E-10 | 2.30E-10 | 1.66E-10 | 5.36E-10 | -2.34E-10 | -6.20E-10 | 3.48E-11 | -6.14E-11 | 1.12E-10 | 0.536306 | 0.345835 |
| NLOT | 3.55E-09 | 5.13E-09 | 3.39E-09 | 5.66E-10 | -2.27E-10 | 7.36E-10 | 7.53E-10 | -1.42E-10 | -5.46E-10 | -8.23E-10 | 2.83E-10 | 3.15E-09 | 0.005413 | 4.212939 |
| COA | 5.75E-10 | -3.10E-10 | -1.02E-09 | -1.06E-09 | -7.57E-10 | -8.75E-10 | -1.26E-09 | 1.13E-09 | 9.62E-10 | 3.73E-10 | 3.00E-10 | -4.34E-10 | 0.538124 | -1.10664 |
| PAA | 2.07E-09 | 4.88E-09 | 3.49E-09 | 1.83E-10 | -8.47E-10 | -6.19E-10 | -1.08E-11 | 4.78E-10 | 3.58E-10 | 3.82E-10 | 2.04E-10 | 2.66E-09 | 0.000059 | 3.623744 |
| TR | -3.93E-10 | 1.23E-09 | -3.46E-10 | -1.12E-09 | 3.94E-10 | -4.27E-11 | -4.08E-10 | 2.27E-10 | -3.66E-10 | -9.75E-11 | 2.89E-10 | -1.56E-10 | 0.397163 | -0.65194 |
| CA1 | 2.78E-10 | 1.58E-09 | 1.67E-09 | -3.28E-10 | -1.44E-10 | -3.92E-10 | -2.46E-10 | 2.42E-10 | 1.77E-10 | 3.57E-10 | -1.74E-11 | 8.03E-10 | 0.001512 | 1.925142 |
| CA2 | -5.02E-11 | 2.48E-09 | 1.72E-09 | -3.56E-10 | -1.44E-10 | -4.33E-10 | -1.73E-12 | 3.82E-10 | -2.17E-10 | 5.70E-10 | -1.54E-10 | 9.50E-10 | 0.015533 | 1.563531 |
| CA3 | -1.97E-10 | 2.19E-09 | 1.71E-09 | -6.75E-10 | -5.48E-10 | -1.12E-09 | -4.08E-10 | 1.44E-09 | -5.27E-10 | 1.17E-09 | -4.26E-11 | 7.62E-10 | 0.101159 | 1.126852 |
| DG | -2.28E-10 | 2.05E-09 | 1.23E-09 | -7.97E-10 | -7.98E-10 | -1.57E-09 | -4.16E-10 | 1.69E-09 | -7.11E-10 | 1.38E-09 | 3.79E-10 | 5.72E-10 | 0.208505 | 0.845263 |
| ENTl | 8.14E-10 | 1.05E-09 | 9.18E-10 | -3.62E-10 | 3.18E-10 | -8.32E-11 | 7.54E-11 | 2.12E-10 | -6.96E-10 | 2.54E-10 | -4.51E-11 | 6.01E-10 | 0.000834 | 2.179513 |
| ENTm | 3.73E-10 | 2.81E-09 | 2.54E-09 | 6.35E-10 | 1.41E-10 | -5.01E-10 | -1.44E-10 | 5.02E-10 | -7.71E-10 | 3.72E-10 | 4.02E-10 | 1.59E-09 | 0.001977 | 3.267499 |
| PAR | 1.21E-09 | 7.30E-10 | 1.20E-09 | 9.16E-10 | -4.65E-10 | -1.78E-10 | 3.59E-10 | 1.91E-11 | -6.58E-11 | -4.40E-11 | 3.59E-10 | 1.01E-09 | 0.001121 | 6.091092 |
| POST | -1.02E-09 | -2.66E-09 | -3.94E-09 | -2.53E-09 | -6.49E-10 | -2.95E-10 | -4.09E-10 | -4.79E-11 | 9.60E-10 | -4.43E-10 | 7.84E-10 | -2.52E-09 | 0.001142 | -5.35711 |
| PRE | 2.35E-09 | 1.39E-10 | 2.96E-10 | 7.98E-10 | -1.07E-09 | -4.80E-10 | -8.24E-11 | 7.95E-10 | 4.14E-10 | 2.57E-10 | 1.08E-10 | 9.05E-10 | 0.100335 | 1.844685 |
| SUB | 4.58E-10 | 2.77E-09 | 3.14E-09 | -4.09E-10 | 2.63E-10 | -8.69E-10 | -5.98E-10 | 2.46E-10 | 1.05E-10 | 9.19E-10 | -8.70E-11 | 1.49E-09 | 0.002402 | 2.042152 |
| ProS | 2.76E-10 | 2.72E-09 | 2.68E-09 | -6.37E-10 | -4.29E-10 | -1.14E-09 | -7.49E-10 | 7.94E-10 | 4.57E-10 | 1.03E-09 | -3.27E-11 | 1.27E-09 | 0.007672 | 1.712154 |
| CLA | 4.71E-09 | 7.79E-09 | 5.39E-09 | 1.57E-09 | 9.36E-10 | 5.13E-10 | 1.28E-09 | -1.09E-09 | -1.63E-09 | -4.12E-10 | 5.24E-10 | 4.85E-09 | 0.000033 | 5.321192 |
| EP | 1.45E-09 | 1.77E-09 | 8.78E-10 | -6.80E-10 | -2.19E-10 | -3.46E-10 | -2.26E-10 | 1.57E-10 | 2.90E-10 | 2.14E-10 | 9.73E-11 | 8.58E-10 | 0.00048 | 1.803964 |
| LA | 2.27E-09 | 2.76E-09 | 9.56E-10 | 1.12E-09 | -8.64E-10 | 4.08E-10 | 7.14E-10 | -2.66E-10 | 6.52E-10 | -3.09E-10 | -3.39E-10 | 1.78E-09 | 0.001599 | 4.623204 |
| BLA | 1.71E-09 | 2.04E-09 | 3.39E-11 | -1.95E-10 | -1.01E-10 | 2.93E-10 | -8.05E-11 | 3.78E-11 | 1.88E-10 | -3.69E-10 | 2.07E-11 | 8.98E-10 | 0.003696 | 1.798042 |
| BMA | 5.40E-10 | -1.01E-10 | -3.69E-09 | -1.48E-09 | -7.11E-10 | -9.26E-10 | -7.48E-10 | 5.31E-10 | 7.35E-10 | 1.92E-10 | 8.11E-10 | -1.17E-09 | 0.146241 | -1.40896 |
| PA | 3.85E-10 | 1.63E-09 | 3.19E-10 | -1.60E-09 | -9.65E-10 | -5.03E-10 | -1.13E-09 | 1.08E-09 | 1.20E-09 | 2.21E-11 | 1.74E-10 | 2.01E-10 | 0.100741 | 0.203883 |
| CP | 2.70E-10 | 1.64E-10 | 6.17E-10 | 1.94E-10 | 2.39E-10 | 3.18E-10 | 2.12E-10 | -2.47E-10 | -1.59E-10 | -1.43E-10 | -1.87E-10 | 3.07E-10 | 0.117926 | 3.571381 |
| STRv | -3.51E-10 | -6.04E-10 | -1.89E-09 | -8.89E-10 | -2.74E-10 | 2.78E-10 | -2.86E-10 | 9.43E-10 | -1.56E-10 | -2.61E-11 | -4.77E-10 | -9.33E-10 | 0.017543 | -3.29398 |
| ACB | 2.70E-10 | 1.75E-09 | 9.92E-10 | -1.28E-10 | 2.18E-10 | 2.02E-10 | 2.83E-10 | -2.58E-10 | -1.43E-10 | 9.02E-11 | -3.53E-10 | 7.15E-10 | 0.01402 | 2.203547 |
| OT | 2.49E-10 | 6.51E-10 | -1.69E-10 | -9.66E-11 | 3.80E-10 | 7.45E-10 | 6.28E-10 | -1.63E-10 | -8.28E-10 | -3.46E-10 | -3.33E-10 | 1.47E-10 | 0.353792 | 0.969377 |
| LSX | -8.40E-11 | 3.19E-09 | 1.06E-09 | 4.67E-10 | -2.82E-10 | 8.52E-10 | 1.87E-09 | -1.06E-09 | -4.27E-10 | -3.00E-10 | -5.49E-10 | 1.14E-09 | 0.339436 | 1.95261 |
| sAMY | 6.67E-11 | -2.26E-10 | -1.70E-09 | -2.18E-09 | -6.14E-11 | -4.30E-10 | 8.42E-11 | -4.25E-10 | 2.17E-10 | 1.31E-10 | 4.60E-10 | -1.01E-09 | 0.722863 | -2.03915 |
| CEA | 1.80E-09 | 5.72E-10 | 6.20E-10 | -2.18E-10 | -7.01E-10 | 1.53E-10 | -1.23E-09 | 9.10E-10 | 1.25E-09 | -6.36E-10 | 1.34E-10 | 7.11E-10 | 0.060635 | 1.723034 |
| MEA | -1.21E-09 | -4.07E-09 | -5.17E-09 | -3.46E-10 | -1.57E-09 | -8.71E-10 | -2.28E-09 | 1.62E-09 | 2.59E-09 | -9.55E-11 | 3.59E-10 | -2.66E-09 | 0.007734 | -2.95653 |
| PAL | -1.44E-09 | -1.45E-10 | 4.40E-10 | -6.51E-10 | 8.48E-11 | 8.93E-10 | -1.85E-10 | 3.76E-10 | 2.45E-11 | -5.74E-10 | -5.96E-10 | -4.52E-10 | 0.473911 | -1.31104 |
| GPe | -4.71E-10 | 2.77E-09 | 4.11E-09 | -4.89E-10 | 2.37E-10 | 3.05E-10 | -6.51E-12 | -6.87E-11 | -1.63E-10 | -2.56E-10 | -3.09E-11 | 1.48E-09 | 0.013081 | 1.350656 |
| GPi | -1.81E-09 | -1.88E-09 | -2.60E-10 | -1.32E-09 | 1.14E-10 | 5.07E-10 | -2.08E-10 | 3.90E-10 | -3.46E-10 | -5.88E-10 | 1.36E-10 | -1.32E-09 | 0.003069 | -3.75438 |
| SI | -3.33E-10 | 2.80E-09 | 1.87E-09 | -5.64E-10 | 1.71E-10 | 8.98E-10 | 7.10E-10 | -1.08E-10 | -7.63E-10 | -5.81E-10 | -2.53E-10 | 9.32E-10 | 0.019294 | 1.193974 |
| MA | -4.28E-10 | 8.19E-10 | -4.28E-09 | -1.00E-09 | 4.74E-10 | 2.26E-10 | 6.55E-10 | -6.05E-11 | -1.35E-09 | -3.95E-11 | 1.69E-10 | -1.23E-09 | 0.879143 | -1.45406 |
| MS | -1.59E-09 | 1.47E-09 | 3.21E-09 | -1.40E-09 | -1.20E-09 | 4.81E-10 | 1.15E-09 | 9.73E-10 | -9.96E-10 | -3.42E-10 | -3.92E-11 | 4.19E-10 | 0.120966 | 0.218838 |
| NDB | -1.17E-09 | 2.80E-09 | -1.20E-10 | -1.33E-09 | 1.11E-09 | 5.16E-10 | 6.50E-10 | -3.85E-10 | -1.71E-09 | -2.11E-10 | 1.43E-10 | 2.76E-11 | 0.320581 | -0.26009 |
| TRS | -3.51E-09 | -6.32E-09 | -7.21E-09 | 2.16E-10 | -2.90E-10 | 1.42E-09 | 1.12E-09 | 2.82E-10 | -8.81E-10 | -7.91E-10 | -7.63E-10 | -4.22E-09 | 0.000032 | -3.16981 |
| BST | -8.82E-10 | 5.79E-09 | 1.02E-09 | -8.98E-10 | -4.95E-10 | 5.10E-10 | 4.79E-11 | 1.17E-09 | -5.44E-10 | -2.20E-10 | -4.52E-10 | 1.25E-09 | 0.564446 | 0.740969 |
| VENT | 5.68E-10 | 2.36E-09 | 2.07E-09 | 1.03E-09 | 6.33E-10 | 1.11E-09 | 2.90E-10 | -6.43E-10 | -1.09E-10 | -7.64E-10 | -4.54E-10 | 1.50E-09 | 0.004142 | 5.247677 |
| SPF | -7.37E-09 | -1.06E-08 | -9.95E-09 | -2.34E-09 | 1.01E-09 | 1.26E-09 | 5.03E-10 | -7.62E-10 | -2.92E-10 | -2.72E-10 | -1.31E-09 | -7.58E-09 | 0.000021 | -5.26528 |
| SPA | -2.50E-09 | -3.11E-09 | 2.18E-09 | 5.74E-10 | -1.36E-09 | 4.60E-10 | 3.34E-09 | -3.12E-09 | 5.12E-10 | -1.24E-09 | 1.43E-09 | -7.17E-10 | 0.398046 | -0.65444 |
| PP | 2.93E-09 | 1.92E-09 | 2.06E-09 | 5.67E-09 | -1.16E-09 | -2.17E-09 | -3.26E-11 | 1.86E-09 | -5.35E-10 | 2.66E-09 | -6.43E-10 | 3.15E-09 | 0.021287 | 3.839814 |
| GENd | -4.12E-10 | 4.77E-09 | 4.93E-09 | -5.16E-10 | 4.05E-10 | 1.34E-09 | -1.27E-10 | 4.27E-11 | 1.51E-11 | -9.43E-10 | -6.84E-10 | 2.18E-09 | 0.011301 | 1.56668 |
| LAT | -8.04E-11 | 6.06E-09 | 2.55E-09 | 2.76E-10 | 8.75E-11 | 4.03E-10 | 2.32E-10 | -2.19E-11 | -1.98E-10 | -1.67E-10 | -3.04E-10 | 2.20E-09 | 0.025954 | 1.86356 |
| ATN | 7.89E-10 | 6.05E-09 | 5.74E-09 | 6.63E-10 | 1.83E-12 | 6.07E-10 | 2.65E-10 | 3.30E-10 | -6.76E-10 | -5.62E-10 | 6.67E-11 | 3.31E-09 | 0.00193 | 2.783173 |

| | | | | | | | | | | | | | | |
|---|---|---|---|---|---|---|---|---|---|---|---|---|---|---|
| **MED** | 7.25E-10 | 8.10E-09 | 5.30E-09 | 5.77E-10 | -9.21E-10 | 7.27E-10 | 1.08E-09 | 5.27E-10 | -5.75E-10 | -4.26E-10 | -3.66E-10 | 3.67E-09 | 0.009219 | 2.549769 |
| **MTN** | 2.75E-09 | 6.26E-09 | 1.39E-09 | 2.56E-09 | -1.87E-09 | -5.33E-10 | 2.40E-09 | 4.62E-10 | -9.89E-10 | 1.10E-09 | -4.96E-10 | 3.23E-09 | 0.263811 | 4.660215 |
| **ILM** | 1.03E-09 | 7.31E-09 | 4.87E-09 | 1.64E-09 | -1.31E-10 | 2.20E-09 | 2.23E-09 | -4.30E-10 | -1.69E-09 | -1.57E-09 | -4.46E-10 | 3.69E-09 | 0.036859 | 3.487259 |
| **RT** | -9.57E-10 | 1.23E-09 | 1.69E-09 | -1.08E-09 | -1.47E-10 | 2.40E-10 | 4.60E-10 | 1.07E-10 | -3.65E-10 | 5.21E-12 | -2.65E-10 | 2.15E-10 | 0.215519 | 0.127795 |
| **GENv** | -1.49E-08 | -2.56E-08 | -1.55E-08 | -4.99E-09 | -3.72E-09 | -2.77E-10 | 2.28E-10 | 9.32E-10 | 3.59E-09 | 1.25E-10 | -1.07E-09 | -1.52E-08 | 0.000011 | -5.14619 |
| **EPI** | -3.94E-09 | -6.27E-09 | -1.04E-08 | -3.56E-09 | 2.96E-10 | 1.05E-09 | -3.69E-10 | 6.80E-10 | -4.27E-10 | -8.15E-10 | -3.89E-10 | -6.05E-09 | 0.000138 | -4.89379 |
| **PVZ** | -2.11E-09 | -1.01E-08 | -1.52E-08 | 1.71E-09 | -3.03E-09 | 2.18E-11 | -2.65E-09 | 4.05E-09 | 3.41E-09 | 5.57E-10 | -2.59E-09 | -6.40E-09 | 0.003399 | -1.97597 |
| **PVR** | -4.12E-09 | -3.29E-09 | -1.21E-08 | -2.33E-10 | 8.97E-11 | -1.37E-10 | -1.70E-09 | 1.03E-09 | 1.85E-09 | 1.30E-09 | -2.46E-09 | -4.94E-09 | 0.002489 | -2.36744 |
| **AHN** | -3.68E-09 | 6.56E-10 | -8.21E-09 | -1.14E-10 | 6.30E-12 | -5.56E-10 | 9.28E-10 | -7.31E-10 | -2.07E-10 | 9.29E-10 | -3.17E-10 | -2.84E-09 | 0.024422 | -1.7984 |
| **MBO** | -3.14E-09 | 3.49E-10 | -1.48E-09 | -1.91E-09 | -1.10E-09 | 1.23E-09 | 4.54E-10 | -7.08E-10 | 1.22E-09 | -2.18E-09 | 1.01E-09 | -1.53E-09 | 0.173349 | -2.32139 |
| **MPN** | -2.35E-10 | 2.61E-09 | -2.24E-09 | 4.94E-09 | -1.15E-09 | 3.88E-10 | -2.19E-09 | 2.51E-09 | 2.29E-09 | 4.64E-10 | -2.41E-09 | 1.28E-09 | 0.64513 | 0.84948 |
| **PMd** | -9.02E-09 | -1.47E-08 | -2.76E-08 | -3.76E-09 | 8.39E-10 | 8.85E-10 | 1.75E-09 | -1.94E-09 | -1.58E-09 | -1.48E-09 | 1.62E-09 | -1.38E-08 | 0.000206 | -3.3241 |
| **PMv** | 4.23E-09 | 4.86E-09 | 9.10E-09 | -2.92E-09 | 1.31E-09 | 1.52E-09 | -2.85E-09 | -1.16E-09 | 2.40E-09 | -3.41E-09 | 1.99E-09 | 3.85E-09 | 0.02919 | 1.815084 |
| **PVHd** | -1.33E-08 | -8.68E-09 | -1.85E-08 | -1.67E-09 | -3.60E-09 | -1.09E-09 | -2.53E-09 | 3.47E-09 | 4.25E-09 | 1.16E-09 | -1.95E-09 | -1.05E-08 | 0.00012 | -3.28027 |
| **VMH** | 8.51E-10 | 3.35E-09 | -3.69E-09 | 2.19E-09 | -2.12E-09 | 1.36E-09 | 1.42E-09 | 1.54E-09 | -1.12E-11 | -3.45E-10 | -1.79E-09 | 6.70E-10 | 0.643191 | 0.455794 |
| **PH** | -1.54E-08 | -2.40E-08 | -3.11E-08 | -8.09E-09 | -2.64E-09 | -1.69E-09 | -7.70E-10 | 1.70E-09 | 2.23E-09 | 1.34E-09 | -3.61E-10 | -1.96E-08 | 0.000006 | -5.15254 |
| **LZ** | -1.64E-09 | -4.76E-10 | -3.77E-09 | -4.40E-10 | 1.15E-10 | 1.19E-09 | 5.73E-10 | -3.66E-10 | -1.80E-10 | -9.33E-10 | -3.45E-10 | -1.59E-09 | 0.007519 | -2.47778 |
| **ZI** | -6.04E-09 | -2.77E-09 | -2.90E-09 | -3.14E-09 | 1.37E-10 | 1.37E-09 | 1.86E-09 | -1.11E-09 | -1.01E-09 | -9.90E-10 | -1.36E-10 | -3.73E-09 | 0.001622 | -3.72759 |
| **MBsen** | 1.30E-09 | 7.90E-10 | 1.53E-09 | 1.44E-09 | -5.34E-10 | -1.82E-10 | 6.49E-11 | -4.01E-10 | 1.35E-09 | 4.78E-10 | -8.00E-10 | 1.27E-09 | 0.008598 | 6.358739 |
| **SCs** | -2.13E-08 | -4.02E-08 | -3.64E-08 | -8.51E-09 | -5.38E-09 | -1.45E-09 | -3.48E-09 | 1.05E-08 | -1.02E-09 | 1.01E-09 | -4.98E-10 | -2.66E-08 | 0.000023 | -5.19715 |
| **IC** | -7.81E-09 | -1.07E-08 | -8.32E-09 | -1.99E-09 | -1.71E-09 | -2.12E-09 | -2.68E-09 | 2.41E-09 | 3.21E-09 | 2.47E-09 | -1.79E-09 | -7.16E-09 | 0.000168 | -4.91818 |
| **MBmot** | -3.28E-09 | -3.40E-09 | -9.73E-10 | -1.69E-09 | 5.96E-10 | 8.07E-10 | 1.27E-10 | -2.70E-10 | -3.30E-10 | -4.69E-10 | -4.03E-10 | -2.34E-09 | 0.003745 | -4.06288 |
| **SNr** | -8.89E-10 | -1.08E-10 | 5.40E-10 | -3.75E-10 | -1.27E-10 | 5.09E-10 | 3.47E-10 | -4.97E-10 | 2.65E-10 | -6.40E-10 | 1.44E-10 | -2.08E-10 | 0.658293 | -0.90126 |
| **VTA** | -1.99E-09 | 2.02E-09 | 1.96E-09 | -2.46E-10 | -5.03E-10 | 1.64E-09 | 1.53E-09 | -4.14E-10 | -5.76E-10 | -1.42E-09 | -1.74E-10 | 4.24E-10 | 0.246478 | 0.208566 |
| **SCm** | -1.06E-08 | -1.04E-08 | -9.23E-09 | -3.56E-09 | -9.05E-10 | -5.05E-10 | -7.22E-11 | -1.41E-09 | 2.84E-09 | 2.25E-10 | -2.80E-10 | -8.42E-09 | <0.000001 | -5.13654 |
| **PAG** | -8.64E-09 | -9.10E-09 | -6.91E-09 | -3.65E-09 | -2.25E-09 | -3.69E-10 | 9.37E-10 | 4.29E-10 | 1.09E-09 | 1.50E-10 | -6.06E-11 | -7.06E-09 | 0.000004 | -5.80774 |
| **PRC** | -2.91E-09 | -7.48E-09 | -9.85E-09 | 1.94E-09 | -9.80E-10 | 1.86E-09 | 5.35E-09 | -4.11E-09 | -5.40E-10 | -1.41E-09 | 4.76E-11 | -4.61E-09 | 0.015861 | -2.11426 |
| **APN** | -5.55E-09 | -2.67E-09 | 9.46E-10 | -1.49E-09 | 7.13E-10 | 2.24E-09 | 1.36E-09 | -8.83E-10 | -1.18E-09 | -1.74E-09 | -3.47E-10 | -2.21E-09 | 0.181775 | -1.70498 |
| **MPT** | -1.17E-08 | -1.96E-08 | -1.72E-08 | -6.42E-09 | -5.44E-09 | 5.65E-09 | 2.88E-09 | -5.05E-10 | 5.93E-09 | -4.20E-09 | -4.35E-09 | -1.37E-08 | 0.000574 | -7.02153 |
| **SNc** | -9.07E-10 | -1.54E-11 | 3.17E-10 | -4.39E-10 | 2.43E-10 | 5.72E-10 | 8.32E-10 | -7.36E-10 | -2.53E-10 | -1.91E-10 | -3.96E-10 | -2.71E-10 | 0.520551 | -1.1966 |
| **PPN** | -1.70E-09 | 1.45E-09 | 1.21E-09 | -1.23E-09 | -4.41E-11 | -5.40E-11 | -1.20E-09 | 6.05E-10 | 9.74E-10 | 1.16E-11 | -3.57E-10 | -6.04E-11 | 0.360881 | -0.37246 |
| **RAmb** | -9.60E-09 | -1.40E-08 | -9.65E-09 | -1.29E-08 | -3.78E-09 | 1.50E-10 | 4.18E-09 | -4.47E-10 | 6.39E-11 | 5.88E-12 | -1.41E-10 | -1.15E-08 | 0.000093 | -15.8591 |
