## Supplementary material for "A developmental brain-wide screen identifies retrosplenial cortex as a key player in the emergence of persistent memory": Table 4

**Table 4. Detailed statistics for whole-brain map P17 mice.**

| Learning Indices | P60 CFC | P60 CFC | P60 CFC | P60 CFC | P60 NS | P60 NS | P60 NS | P60 NS | P60 NS | P60 NS | P60 NS | Difference CFC-NS | p value (T-test) | Salience |
|---|---|---|---|---|---|---|---|---|---|---|---|---|---|---|
| **FRP** | 1.17E-09 | 1.38E-09 | 4.86E-10 | 4.07E-10 | -1.93E-09 | -1.44E-09 | -2.52E-10 | -1.56E-09 | 5.69E-09 | 4.63E-11 | -7.58E-10 | 8.90E-10 | 0.527332 | 3.651462 |
| **MOp** | 2.23E-09 | -7.54E-10 | -1.04E-09 | -6.70E-10 | 1.43E-11 | -6.32E-10 | 1.33E-09 | -6.49E-11 | -1.68E-10 | -5.74E-10 | -4.92E-11 | -3.76E-11 | 0.954916 | 0.23153 |
| **MOs** | 1.27E-09 | -3.32E-11 | -2.73E-10 | 6.91E-11 | -2.70E-10 | -2.65E-10 | 1.45E-09 | -3.92E-10 | -3.59E-10 | -1.47E-10 | -8.10E-11 | 2.66E-10 | 0.538612 | 0.990953 |
| **SSp** | 2.22E-09 | -3.58E-10 | -5.14E-10 | -3.24E-11 | -3.29E-10 | -2.91E-10 | 1.36E-09 | -1.18E-10 | -3.01E-10 | -4.21E-10 | -3.06E-12 | 3.44E-10 | 0.555456 | 0.784602 |
| **SSs** | 2.00E-09 | 5.81E-10 | -6.05E-10 | 1.10E-10 | -9.03E-12 | -3.95E-10 | 1.26E-09 | -2.11E-10 | -3.88E-10 | -2.65E-10 | -7.96E-11 | 5.34E-10 | 0.310714 | 1.182102 |
| **GU** | 2.39E-09 | 1.57E-09 | 1.00E-09 | 1.86E-09 | 1.18E-09 | -6.67E-10 | 1.67E-09 | -2.30E-10 | -1.73E-09 | 9.21E-11 | -3.61E-10 | 1.71E-09 | 0.022043 | 4.28719 |
| **VISC** | 2.11E-09 | 1.39E-09 | -1.09E-10 | 6.63E-10 | 7.61E-10 | -6.29E-10 | 1.56E-09 | -3.52E-10 | -1.14E-09 | -2.57E-11 | -2.34E-10 | 1.02E-09 | 0.109779 | 2.21783 |
| **AUD** | 1.33E-09 | 3.47E-10 | 1.35E-09 | 3.75E-10 | 2.49E-10 | -6.17E-10 | 1.59E-09 | 2.59E-10 | -9.05E-10 | -1.03E-09 | 2.65E-10 | 8.77E-10 | 0.118688 | 3.057732 |
| **VIS** | 7.88E-10 | 1.68E-09 | 4.46E-09 | 2.08E-09 | 7.58E-10 | -3.10E-11 | -9.80E-10 | -3.00E-10 | 1.19E-10 | 4.33E-10 | 9.34E-11 | 2.24E-09 | 0.006409 | 4.215472 |
| **ACA** | -2.01E-09 | -3.59E-11 | 2.51E-09 | 1.66E-09 | -8.68E-11 | -7.76E-11 | 1.98E-09 | -8.67E-10 | -9.99E-10 | 1.19E-09 | -1.06E-09 | 5.20E-10 | 0.593508 | 0.304896 |
| **PL** | -9.29E-10 | 9.64E-10 | 2.49E-09 | 1.33E-09 | -5.66E-10 | 5.03E-10 | 1.54E-09 | -9.50E-10 | -9.74E-10 | 1.14E-09 | -5.41E-10 | 9.43E-10 | 0.231952 | 1.262661 |
| **ILA** | -1.60E-09 | 8.37E-10 | 1.82E-09 | 3.12E-09 | 6.09E-11 | 6.33E-10 | 1.42E-09 | -2.12E-11 | -2.22E-09 | 1.12E-09 | -8.45E-10 | 1.02E-09 | 0.315361 | 0.878819 |
| **ORB** | -1.68E-10 | 1.62E-09 | 1.60E-09 | 1.48E-09 | -6.76E-10 | 1.19E-10 | 1.35E-09 | -1.91E-10 | -4.62E-10 | -3.35E-11 | -1.35E-10 | 1.14E-09 | 0.03502 | 2.8109 |
| **AI** | 9.11E-10 | 1.27E-09 | -1.99E-10 | -9.90E-11 | 1.71E-10 | -1.71E-10 | 1.16E-09 | -1.70E-10 | -9.13E-10 | -1.75E-10 | 5.58E-11 | 4.76E-10 | 0.279218 | 1.575702 |
| **RSP** | 6.27E-10 | 1.08E-09 | 9.00E-10 | 2.32E-09 | 3.67E-10 | 2.79E-10 | 3.57E-10 | -3.53E-11 | -1.17E-09 | 4.61E-10 | -1.72E-10 | 1.22E-09 | 0.013535 | 3.608575 |
| **PTLp** | -8.45E-10 | -3.34E-10 | 1.68E-09 | 1.70E-09 | 7.88E-10 | 6.82E-10 | -9.50E-10 | 7.20E-10 | -1.52E-09 | 4.54E-10 | -3.18E-11 | 5.30E-10 | 0.450539 | 0.74585 |
| **TEa** | 2.31E-09 | 5.41E-10 | 2.27E-09 | 3.46E-10 | -2.34E-11 | -5.11E-10 | 7.82E-10 | 8.13E-10 | -1.70E-10 | -6.72E-10 | -3.88E-10 | 1.39E-09 | 0.020191 | 2.652967 |
| **PERI** | 3.97E-09 | 2.87E-09 | 2.36E-09 | 8.64E-10 | 2.89E-10 | -1.17E-09 | 8.92E-12 | -1.17E-09 | 2.10E-09 | 3.90E-10 | -5.07E-10 | 2.52E-09 | 0.007891 | 3.38685 |
| **ECT** | 1.11E-09 | 1.48E-09 | 1.42E-09 | 7.45E-10 | -5.48E-10 | 2.12E-10 | 1.43E-09 | -2.03E-10 | -7.48E-10 | 6.04E-10 | -7.09E-10 | 1.19E-09 | 0.021626 | 9.638945 |
| **TT** | -8.25E-10 | 1.75E-09 | -6.89E-10 | 4.34E-10 | 2.18E-10 | 5.94E-10 | -1.15E-10 | -5.39E-10 | -9.25E-10 | 5.06E-10 | 4.17E-10 | 1.44E-10 | 0.788816 | 0.139493 |
| **DP** | -1.50E-09 | 7.55E-10 | 2.34E-10 | 6.61E-10 | -2.35E-10 | 5.46E-10 | 1.06E-09 | -1.09E-09 | -9.80E-10 | 1.30E-09 | -4.10E-10 | 8.75E-12 | 0.989055 | -0.20978 |
| **PIR** | -1.15E-09 | -1.79E-10 | -2.01E-09 | -1.58E-09 | 1.08E-10 | -1.80E-10 | -1.16E-09 | -1.85E-10 | 1.15E-09 | -6.33E-10 | 8.60E-10 | -1.22E-09 | 0.036897 | -3.84192 |
| **NLOT** | -2.65E-10 | 2.01E-09 | 5.08E-09 | 4.98E-09 | -8.95E-10 | -2.75E-10 | -6.63E-10 | -3.20E-09 | 3.77E-09 | 2.89E-09 | -1.33E-09 | 2.91E-09 | 0.094834 | 2.513524 |
| **COA** | -8.39E-11 | 2.14E-09 | 2.39E-09 | -4.09E-10 | 3.42E-10 | 6.11E-10 | -2.32E-09 | 2.51E-10 | 5.22E-10 | 3.53E-10 | 3.95E-10 | 9.87E-10 | 0.220653 | 1.517139 |
| **PAA** | 1.15E-09 | 2.75E-09 | 1.88E-09 | 2.05E-10 | 5.31E-10 | 6.21E-10 | -1.81E-09 | -9.68E-10 | 2.73E-10 | 5.94E-10 | 9.81E-10 | 1.47E-09 | 0.0514 | 3.62945 |
| **TR** | 6.34E-10 | 1.58E-09 | 1.05E-09 | -6.96E-11 | 6.16E-10 | -3.63E-10 | -4.20E-10 | -5.27E-11 | 3.49E-10 | 7.97E-10 | -8.79E-10 | 7.92E-10 | 0.079201 | 2.887428 |
| **CA1** | -9.09E-10 | -9.99E-11 | -9.35E-10 | -1.05E-09 | 4.20E-11 | 6.78E-11 | -3.40E-10 | -2.43E-10 | 3.07E-10 | 5.95E-10 | -3.57E-10 | -7.58E-10 | 0.011738 | -3.54911 |
| **CA2** | -1.98E-09 | -2.63E-10 | -2.06E-09 | -2.54E-09 | -5.82E-11 | -5.93E-10 | -4.81E-10 | 6.05E-11 | 1.46E-09 | -1.65E-10 | -3.17E-10 | -1.70E-09 | 0.00834 | -3.58239 |
| **CA3** | -2.24E-09 | -1.81E-09 | -4.33E-09 | -4.59E-09 | -1.19E-10 | 4.07E-10 | -6.06E-10 | -1.53E-10 | 2.04E-10 | 1.05E-09 | -6.35E-10 | -3.26E-09 | 0.000402 | -5.46624 |
| **DG** | -1.45E-09 | -7.56E-10 | -3.88E-09 | -4.26E-09 | -2.01E-10 | 2.88E-10 | -6.55E-10 | -2.54E-10 | 5.83E-10 | 9.98E-10 | -6.33E-10 | -2.60E-09 | 0.005072 | -3.56189 |
| **ENTl** | -1.61E-10 | 8.81E-10 | 2.97E-10 | 1.96E-10 | -1.52E-10 | -2.01E-10 | 3.62E-10 | -9.77E-11 | 3.06E-10 | 3.13E-10 | -5.42E-10 | 3.05E-10 | 0.22442 | 1.392592 |
| **ENTm** | -6.50E-10 | 3.34E-10 | -1.82E-10 | 2.61E-10 | 4.74E-10 | -7.04E-10 | 5.49E-10 | -8.96E-10 | 5.11E-10 | 4.97E-10 | -4.35E-10 | -5.90E-11 | 0.876893 | -0.49 |
| **PAR** | -3.54E-10 | 6.97E-10 | -1.12E-09 | -2.60E-10 | -1.98E-10 | -5.61E-10 | 8.46E-10 | -6.78E-10 | 7.93E-10 | 5.32E-10 | -7.61E-10 | -2.55E-10 | 0.586249 | -0.9417 |
| **POST** | 8.70E-10 | 2.29E-09 | -6.20E-10 | 2.72E-09 | 9.56E-10 | 1.51E-10 | -2.38E-09 | 5.39E-10 | 4.42E-10 | 1.41E-10 | 2.38E-10 | 1.30E-09 | 0.130467 | 2.210986 |
| **PRE** | 1.19E-09 | -1.02E-11 | -1.53E-10 | 3.84E-10 | -6.04E-10 | -3.45E-10 | -7.79E-10 | -1.29E-10 | 2.08E-09 | -8.80E-11 | -1.91E-10 | 3.61E-10 | 0.516848 | 1.373999 |
| **SUB** | -4.05E-10 | 4.53E-10 | -1.88E-10 | 6.88E-10 | -3.83E-10 | 4.75E-10 | -9.07E-10 | -4.73E-10 | 8.09E-10 | 1.11E-09 | -4.65E-10 | 1.13E-10 | 0.799684 | 0.425313 |
| **ProS** | -1.62E-09 | 3.43E-10 | -1.86E-09 | -1.32E-09 | 2.60E-10 | 4.64E-10 | -7.87E-10 | -5.34E-10 | -5.05E-11 | 1.27E-09 | -4.15E-10 | -1.14E-09 | 0.051359 | -2.57132 |
| **CLA** | -1.74E-10 | 1.72E-09 | 5.00E-10 | -2.03E-10 | -1.09E-09 | 9.38E-10 | -1.11E-10 | 1.10E-09 | -5.37E-10 | -2.32E-10 | -5.26E-11 | 4.58E-10 | 0.395957 | 1.011057 |
| **EP** | -9.94E-10 | 6.72E-10 | 1.88E-10 | 6.16E-10 | -9.71E-11 | 2.00E-10 | 1.31E-10 | 9.79E-11 | -3.63E-10 | 2.39E-11 | 2.55E-11 | 1.18E-10 | 0.698518 | 0.038167 |
| **LA** | 1.60E-09 | 1.42E-09 | 2.88E-09 | 1.14E-09 | 2.93E-10 | -1.54E-11 | 2.66E-11 | -1.94E-11 | -3.93E-10 | -1.89E-10 | 2.96E-10 | 1.76E-09 | 0.000282 | 6.788331 |
| **BLA** | 2.27E-10 | 1.48E-09 | 5.63E-10 | 7.07E-10 | 6.16E-10 | 4.57E-11 | -2.52E-12 | -5.49E-11 | -7.97E-10 | 2.82E-10 | -3.10E-11 | 7.37E-10 | 0.032418 | 3.53315 |
| **BMA** | -1.44E-09 | 6.18E-10 | 1.07E-11 | -7.81E-10 | 5.31E-10 | 6.69E-10 | -1.60E-09 | -1.12E-09 | 4.86E-11 | 4.91E-10 | 1.21E-09 | -4.32E-10 | 0.501068 | -1.16611 |
| **PA** | -4.78E-10 | 1.39E-09 | 8.46E-11 | 1.49E-10 | -5.26E-10 | 6.13E-10 | -9.49E-10 | 1.96E-10 | 5.06E-10 | 7.73E-10 | -4.92E-10 | 2.69E-10 | 0.561246 | 0.584284 |
| **CP** | -5.20E-10 | -5.92E-10 | -9.95E-10 | -9.93E-10 | 1.74E-10 | 1.36E-11 | -2.09E-10 | -6.97E-11 | 1.40E-11 | 3.29E-10 | -2.12E-10 | -7.81E-10 | 0.000285 | -7.92695 |
| **STRv** | -5.32E-10 | -1.07E-09 | -1.11E-09 | 5.10E-10 | 6.51E-10 | 4.48E-11 | -1.29E-09 | -2.98E-10 | 2.42E-10 | -5.17E-10 | 1.20E-09 | -5.57E-10 | 0.290809 | -1.71484 |
| **ACB** | -1.73E-10 | 6.38E-10 | 4.83E-10 | 2.03E-10 | -1.04E-10 | 7.55E-11 | 9.25E-11 | 1.55E-10 | -1.91E-10 | -3.74E-10 | 3.15E-10 | 2.92E-10 | 0.13043 | 1.571225 |
| **OT** | -4.70E-11 | 1.41E-09 | 5.13E-10 | 4.69E-10 | -2.29E-11 | 3.34E-10 | -6.09E-10 | -2.01E-10 | 1.87E-10 | 7.98E-10 | -3.65E-10 | 5.68E-10 | 0.114372 | 2.166769 |
| **LSX** | -1.03E-09 | -1.37E-09 | 2.50E-09 | 9.74E-10 | -9.89E-10 | 8.36E-10 | 6.17E-10 | 7.38E-10 | -9.01E-10 | 4.42E-10 | -6.90E-10 | 2.60E-10 | 0.746021 | 0.191213 |
| **sAMY** | -2.08E-10 | 8.20E-10 | 2.31E-09 | 2.34E-09 | -4.41E-10 | 2.53E-10 | -1.38E-09 | -1.83E-09 | 2.09E-09 | 6.51E-10 | 8.36E-10 | 1.29E-09 | 0.151692 | 2.282984 |
| **CEA** | -5.09E-10 | -2.60E-10 | 2.67E-10 | 1.77E-09 | 1.47E-09 | -2.60E-10 | -5.60E-10 | 5.68E-10 | -1.11E-09 | 1.28E-10 | -2.08E-10 | 3.15E-10 | 0.590719 | 0.613912 |
| **MEA** | 7.19E-10 | 5.32E-09 | 7.77E-09 | 3.54E-09 | -6.43E-10 | 2.51E-10 | -1.87E-09 | -1.23E-09 | 2.57E-09 | 6.34E-10 | 4.43E-10 | 4.32E-09 | 0.009219 | 4.000079 |
| **PAL** | -2.99E-10 | -2.10E-10 | -5.95E-10 | -6.47E-10 | 7.06E-12 | -2.12E-10 | -1.17E-10 | 3.28E-10 | 3.56E-10 | 8.47E-11 | -4.82E-10 | -4.33E-10 | 0.03247 | -4.82866 |
| **GPe** | -1.72E-09 | -2.01E-09 | -3.15E-09 | -2.09E-09 | 1.46E-10 | 2.47E-10 | -1.00E-10 | -1.34E-10 | -3.73E-10 | 6.46E-10 | -3.37E-10 | -2.26E-09 | 0.000029 | -14.7407 |
| **GPi** | -2.26E-09 | -3.08E-09 | -3.96E-09 | -3.65E-09 | -6.28E-10 | -3.12E-10 | 1.56E-09 | 8.64E-10 | -4.23E-10 | -5.72E-10 | -6.67E-10 | -3.21E-09 | 0.000172 | -15.0606 |
| **SI** | -6.10E-12 | 4.35E-10 | -4.31E-11 | 2.41E-10 | 5.49E-11 | 2.29E-10 | -4.40E-10 | -3.02E-10 | 6.42E-11 | 2.20E-10 | 2.46E-10 | 1.46E-10 | 0.390261 | 1.623531 |
| **MA** | -8.73E-10 | -1.93E-11 | -1.46E-09 | -9.16E-10 | -3.70E-11 | 1.59E-10 | -2.96E-10 | -1.06E-09 | 5.04E-10 | 6.18E-10 | 2.37E-10 | -8.36E-10 | 0.046525 | -3.39034 |
| **MS** | 9.52E-10 | 2.34E-09 | 1.66E-09 | 3.67E-09 | 4.14E-10 | 1.10E-09 | -6.94E-10 | -8.46E-10 | -1.28E-09 | 1.39E-09 | 2.40E-10 | 2.11E-09 | 0.011511 | 4.154838 |
| **NDB** | 1.60E-09 | 2.20E-09 | 4.04E-10 | 1.22E-09 | -9.74E-10 | 3.40E-10 | -5.87E-10 | -7.44E-10 | 1.46E-09 | 1.76E-10 | 3.86E-10 | 1.35E-09 | 0.026973 | 4.054782 |
| **TRS** | 1.21E-10 | 2.43E-09 | 3.55E-09 | 9.91E-10 | -7.67E-10 | 2.22E-10 | 3.59E-10 | 7.91E-10 | 1.75E-10 | 1.07E-09 | -1.82E-09 | 1.77E-09 | 0.042337 | 3.000081 |
| **BST** | -6.04E-10 | 5.44E-10 | 4.22E-09 | 3.31E-09 | -6.23E-10 | 4.63E-10 | -2.02E-09 | 1.70E-09 | 1.02E-09 | -9.65E-10 | 3.34E-10 | 1.88E-09 | 0.106443 | 1.780476 |
| **VENT** | 3.51E-10 | -9.54E-10 | -5.20E-10 | -3.02E-10 | 1.71E-10 | -9.43E-11 | 2.87E-10 | 4.13E-10 | -4.15E-10 | 2.36E-10 | -6.09E-10 | -3.55E-10 | 0.234838 | -1.1631 |
| **SPF** | 1.45E-09 | -4.15E-09 | -1.74E-09 | -2.51E-09 | 2.25E-09 | -1.39E-10 | -2.71E-09 | 1.40E-10 | -1.66E-10 | 8.35E-10 | -2.21E-11 | -1.76E-09 | 0.156294 | -1.35186 |
| **SPA** | -5.19E-10 | 1.75E-09 | 2.57E-09 | 2.58E-09 | -4.68E-11 | -1.17E-09 | 4.39E-09 | 1.30E-10 | -2.06E-09 | -1.42E-09 | -1.73E-10 | 1.65E-09 | 0.205919 | 2.219584 |
| **PP** | 9.86E-09 | 3.08E-09 | 7.50E-10 | 2.12E-09 | 5.63E-10 | -4.65E-10 | 1.28E-09 | -1.30E-09 | -4.60E-10 | 7.83E-10 | -3.33E-10 | 3.94E-09 | 0.030384 | 1.915405 |
| **GENd** | -1.18E-09 | -1.44E-09 | -2.99E-09 | -2.14E-09 | -2.13E-10 | -2.07E-10 | 9.89E-10 | 6.97E-10 | -5.69E-10 | -5.29E-10 | -2.96E-10 | -1.92E-09 | 0.001545 | -7.57229 |
| **LAT** | -6.45E-10 | -1.70E-09 | -1.53E-09 | -1.10E-09 | 3.72E-10 | -2.15E-10 | 3.36E-10 | -4.36E-11 | -4.05E-10 | -1.22E-10 | 5.27E-11 | -1.24E-09 | 0.000366 | -9.90992 |
| **ATN** | -2.25E-10 | -1.48E-09 | -3.75E-10 | 7.86E-10 | -1.77E-10 | -2.64E-10 | 8.35E-10 | -3.23E-10 | 4.67E-11 | -8.05E-11 | -8.46E-11 | -3.17E-10 | 0.437987 | -0.71612 |

| | | | | | | | | | | | | | |
|---|---|---|---|---|---|---|---|---|---|---|---|---|---|
| **MED** | -5.14E-10 | -1.52E-09 | 3.27E-10 | -6.81E-10 | -6.15E-10 | -4.31E-10 | 1.53E-09 | -3.24E-10 | 1.35E-10 | -9.99E-10 | 5.33E-10 | -5.74E-10 | 0.292819 | -2.08732 |
| **MTN** | -1.19E-09 | -2.61E-09 | 2.61E-09 | 6.63E-11 | 2.70E-10 | -1.30E-09 | 3.32E-09 | -1.39E-09 | -8.18E-10 | -1.80E-09 | 1.45E-09 | -2.39E-10 | 0.851778 | -0.45588 |
| **ILM** | -2.31E-10 | -1.80E-09 | -4.33E-10 | 7.82E-10 | 3.50E-10 | -4.65E-10 | 2.13E-09 | -9.56E-10 | -1.19E-09 | -7.17E-11 | 1.65E-10 | -4.15E-10 | 0.557002 | -0.82736 |
| **RT** | -1.59E-09 | -2.51E-09 | -2.75E-09 | -2.83E-09 | 1.53E-10 | -2.71E-12 | -9.91E-11 | 9.38E-10 | -3.94E-10 | 2.40E-10 | -8.55E-10 | -2.42E-09 | 0.000073 | -13.7363 |
| **GENv** | -2.07E-09 | -4.60E-09 | -1.52E-09 | -6.01E-09 | 6.68E-11 | -1.06E-09 | 8.31E-10 | 3.28E-09 | -4.45E-10 | -1.85E-09 | -1.27E-09 | -3.49E-09 | 0.015337 | -3.89309 |
| **EPI** | -2.81E-09 | -2.08E-09 | -4.74E-09 | -2.50E-09 | -8.41E-10 | 2.02E-10 | -2.56E-09 | 4.34E-10 | 3.01E-09 | 1.16E-09 | -1.30E-09 | -3.04E-09 | 0.015173 | -7.14145 |
| **PVZ** | -1.45E-09 | 3.09E-09 | 3.56E-09 | 6.00E-09 | 8.40E-10 | -5.57E-10 | -6.76E-09 | 4.69E-09 | 3.90E-09 | -1.66E-09 | -7.32E-10 | 2.84E-09 | 0.23936 | 1.723141 |
| **PVR** | 5.02E-10 | 6.77E-10 | 5.90E-10 | 3.99E-09 | -1.17E-09 | -2.29E-10 | -1.32E-09 | 2.74E-09 | 1.57E-09 | -3.27E-09 | 1.25E-09 | 1.50E-09 | 0.249224 | 1.91163 |
| **AHN** | 5.12E-10 | -2.09E-09 | 1.30E-09 | 2.10E-09 | -1.51E-09 | 1.02E-09 | 3.86E-10 | 3.16E-10 | -3.87E-10 | -7.70E-10 | 9.38E-10 | 4.56E-10 | 0.587986 | 0.704523 |
| **MBO** | -1.87E-09 | -6.19E-10 | -4.74E-09 | -1.08E-09 | -4.06E-11 | -1.22E-09 | 1.37E-09 | 3.49E-10 | 5.37E-10 | -1.76E-09 | 4.32E-10 | -2.03E-09 | 0.044336 | -3.13665 |
| **MPN** | 1.46E-09 | 3.30E-09 | 1.02E-08 | 1.16E-08 | -2.20E-09 | -2.27E-10 | 9.47E-10 | 1.89E-09 | 1.38E-09 | -2.04E-09 | -1.30E-10 | 6.69E-09 | 0.008315 | 3.048977 |
| **PMd** | 6.59E-09 | -5.03E-09 | -1.50E-09 | 1.03E-09 | -4.43E-10 | -2.42E-09 | 3.82E-09 | -2.33E-09 | 2.09E-09 | -5.48E-10 | -5.10E-10 | 3.21E-10 | 0.882881 | 0.427767 |
| **PMv** | 9.88E-10 | -4.16E-11 | 3.43E-09 | 4.41E-09 | -2.55E-09 | -1.35E-10 | 1.87E-09 | 4.14E-10 | 1.44E-09 | -1.55E-09 | 2.03E-10 | 2.24E-09 | 0.071831 | 2.485204 |
| **PVHd** | 3.93E-09 | 5.99E-09 | -5.62E-09 | 2.83E-10 | -1.82E-09 | 2.81E-10 | -5.55E-10 | 2.03E-09 | 8.42E-10 | -4.93E-09 | 3.66E-09 | 1.22E-09 | 0.613479 | 0.728883 |
| **VMH** | 2.96E-09 | 3.82E-10 | 3.99E-09 | 3.58E-09 | -7.28E-10 | 3.19E-10 | 6.48E-10 | -6.98E-10 | 1.45E-10 | 5.47E-10 | -1.60E-10 | 2.72E-09 | 0.002438 | 3.784324 |
| **PH** | -3.30E-09 | -3.11E-09 | -7.48E-09 | -2.55E-11 | -1.52E-09 | 4.77E-10 | -4.47E-09 | 4.77E-09 | 2.50E-09 | -5.51E-09 | 3.18E-09 | -3.40E-09 | 0.171853 | -3.05679 |
| **LZ** | 1.12E-09 | -9.68E-10 | -4.25E-10 | 2.32E-09 | -2.37E-10 | 6.48E-11 | -6.43E-10 | 5.20E-11 | 6.35E-10 | -6.42E-10 | 7.29E-10 | 5.16E-10 | 0.419213 | 1.004992 |
| **ZI** | -5.25E-10 | -2.24E-09 | -1.66E-09 | 1.89E-10 | 1.00E-11 | 3.21E-10 | -4.02E-10 | -3.42E-10 | 2.41E-11 | 7.22E-10 | -2.13E-10 | -1.07E-09 | 0.038948 | -2.37752 |
| **MBsen** | 1.12E-09 | -6.76E-11 | 2.44E-11 | 2.93E-09 | 6.17E-11 | -7.10E-10 | 6.28E-10 | -6.78E-10 | 5.48E-10 | -1.76E-09 | 1.71E-09 | 1.03E-09 | 0.212598 | 1.764861 |
| **SCs** | -6.24E-09 | 6.73E-09 | 7.20E-09 | -1.61E-10 | 8.62E-10 | 2.50E-09 | -7.26E-09 | 3.91E-09 | -6.61E-10 | -1.67E-09 | 2.50E-09 | 1.86E-09 | 0.551413 | 0.297031 |
| **IC** | -6.24E-09 | -5.79E-09 | -2.49E-09 | -4.07E-09 | -5.73E-10 | 9.85E-10 | -3.25E-09 | 3.36E-09 | 2.23E-10 | -3.15E-09 | 2.18E-09 | -4.61E-09 | 0.010287 | -4.48708 |
| **MBmot** | -9.85E-10 | -1.26E-09 | -8.93E-10 | 1.03E-11 | -7.81E-11 | 3.39E-10 | -6.90E-10 | -4.59E-11 | 1.84E-10 | 2.85E-10 | 8.20E-11 | -7.93E-10 | 0.015642 | -3.19589 |
| **SNr** | -2.37E-10 | 5.91E-10 | -1.10E-09 | -1.08E-09 | 7.57E-11 | 1.63E-12 | -2.42E-10 | 8.65E-11 | 9.16E-11 | 5.77E-11 | -6.34E-11 | -4.58E-10 | 0.158801 | -1.41413 |
| **VTA** | 1.71E-09 | 9.34E-10 | 2.10E-10 | 9.51E-10 | -8.14E-10 | 3.41E-10 | -8.31E-11 | 9.19E-10 | 1.25E-10 | -5.03E-10 | -5.62E-11 | 9.61E-10 | 0.026728 | 2.839845 |
| **SCm** | -3.94E-09 | -3.47E-09 | -2.21E-09 | -1.63E-09 | -3.21E-12 | 8.45E-10 | -3.89E-09 | 5.94E-10 | 1.77E-09 | 4.71E-11 | 7.95E-10 | -2.83E-09 | 0.02068 | -4.27968 |
| **PAG** | -3.71E-09 | -2.56E-09 | -7.03E-10 | 1.40E-10 | -1.64E-10 | 1.05E-09 | -4.00E-09 | 1.25E-09 | 1.47E-09 | -6.57E-11 | 5.99E-10 | -1.73E-09 | 0.16787 | -1.98659 |
| **PRC** | 7.13E-10 | 1.14E-09 | 5.17E-09 | 7.07E-09 | -2.05E-09 | -3.98E-10 | -2.71E-09 | -1.24E-09 | 5.83E-09 | -1.78E-09 | 2.18E-09 | 3.55E-09 | 0.096845 | 2.544386 |
| **APN** | 3.08E-10 | -2.94E-09 | -1.07E-09 | 1.05E-09 | 4.95E-10 | 9.62E-10 | -1.74E-09 | -1.05E-09 | -5.45E-11 | 2.09E-09 | -3.12E-10 | -7.19E-10 | 0.449816 | -0.68715 |
| **MPT** | 4.68E-09 | 4.64E-10 | 5.08E-09 | -6.47E-09 | 5.52E-10 | 4.48E-09 | -3.80E-09 | -2.02E-09 | -4.82E-09 | -3.01E-09 | 9.18E-09 | 8.57E-10 | 0.797938 | 0.433914 |
| **SNc** | -2.78E-11 | 7.51E-10 | -7.91E-10 | 2.56E-10 | 7.76E-11 | -2.34E-10 | 7.24E-10 | -2.57E-10 | -2.42E-10 | 3.50E-12 | -9.92E-11 | 5.09E-11 | 0.866044 | 0.21856 |
| **PPN** | -2.01E-09 | -7.62E-10 | -1.05E-10 | 1.94E-10 | 1.33E-09 | 4.66E-10 | -2.16E-09 | 3.69E-10 | -5.59E-10 | 1.06E-09 | -2.81E-10 | -7.02E-10 | 0.341025 | -1.46242 |
| **RAmb** | -6.82E-09 | -6.91E-09 | -4.98E-09 | -7.58E-09 | -1.41E-09 | 1.71E-09 | -2.04E-09 | 3.24E-09 | -7.27E-10 | -1.61E-09 | 7.67E-10 | -6.56E-09 | 0.000192 | -7.37465 |
